# Real-time conversion of tissue-scale mechanical forces into an interdigitated growth pattern

**DOI:** 10.1101/2021.02.06.428645

**Authors:** Samuel A. Belteton, Wenlong Li, Makoto Yanagisawa, Faezeh A. Hatam, Madeline I. Quinn, Margarete K. Szymanski, Mathew W. Marley, Joseph A. Turner, Daniel B. Szymanski

**Affiliations:** Department of Botany and Plant Pathology, Purdue University, West Lafayette, IN 47907; Department of Mechanical and Materials Engineering, University of Nebraska-Lincoln, Lincoln, NE 40247; Department of Botany, University of Wisconsin, Madison, Wisconsin 53706; Department of Biochemistry, Indiana University, Bloomington, IN 47405; Department of Biological Sciences, Purdue University, West Lafayette, IN 47907

**Author notes:** Corresponding Author: Daniel B.

## Abstract

The leaf epidermis is a dynamic biomechanical shell that integrates growth across spatial scales to influence organ morphology. Pavement cells, the fundamental unit of this tissue, morph irreversibly into highly lobed cells that drive planar leaf expansion. Here we define how tissue-scale cell wall tensile forces and the microtubule-cellulose synthase systems pattern interdigitated growth in real-time. A morphologically potent subset of cortical microtubules span the periclinal and anticlinal cell faces to pattern cellulose fibers that generate a patch of anisotropic wall. The result is local polarized growth that is mechanically coupled to the adjacent cell via a pectin-rich middle lamella, and this drives lobe formation. Finite element pavement cell models revealed cell wall tensile stress as an upstream patterning element that links cell- and tissue-scale biomechanical parameters to interdigitated growth. Cell lobing in leaves is evolutionarily conserved, occurs in multiple cell types, and is associated with important agronomic traits. Our general mechanistic models of lobe formation provide a foundation to analyze the cellular basis of leaf morphology and function.

## Introduction

Leaves are specialized planar organs optimized for photosynthesis and gas exchange. Their anatomy varies wildly and reflects complex interactions between the environment and lifecycle^1^. In agronomic settings, leaf traits track with yield, and at a cellular scale, the morphology of lobed mesophyll cells is a target for crop improvement^2^. The epidermis functions as a biomechanical shell that controls the growth properties at organ-scales^3-5^. The sizes and shapes of leaves are ultimately determined by the slow, irreversible expansion of pavement cells that can increase in volume 100-fold^6,7^. A lobed pavement cell morphology is an evolutionarily conserved feature^8^, and this indigitated growth mode drives organ expansion and likely increases the mechanical toughness to large thin leaves. Underlying mesophyll cells form lobes in a cell autonomous manner, and this is one of many important cellular traits that underlie more efficient CO_2_ transfer for photosynthesis and is correlated with increased yield^9^. Despite the widespread occurrence and broad importance of lobed morphogenesis, the mechanisms by which polarized growth process occurs is a long-standing and controversial research question^10-15^.

The patterns of lobed growth are determined both by the mechanical properties of the tough outer cell wall and the tissue context in which they develop. Plant cells grow in a symplastic manner; adjacent cell walls are glued to each other via a pectin-rich middle lamella^16^. Growth rates of the two cells are equal at the cell-cell interface, but subcellular growth rate gradients are expected because adjacent cells can differ in areal growth rates by more than a factor of two^17-19^. Tensile forces in the cell wall are strongly influenced by the tissue context. The epidermis tissue is under tension due to the pressure exerted by underlying cells. The outer periclinal (parallel to the leaf surface) cell wall bears more turgor-generated cell wall tensile force than the adherent anticlinal (perpendicular to the leaf surface) wall because the outer wall is not physically coupled to neighboring cells (Figure 1a). A combination of pavement cell finite element (FE) modeling and microtubule imaging showed examples in which patterns of tensile stress and microtubules were similar^20^. However, neither a quantitative analysis of MTs and stress nor an analysis of cell shape change were performed, leaving open the question of how cell wall stresses might influence growth patterns in this cell type. Lobe formation also occurs in a more cell autonomous manner in partially or non-adherent leaf mesophyll cells in many plant species^11^, creating a labyrinth of intercellular airspace that increases the rate of CO_2_ transfer for photosynthesis^2^.

**Figure 1.**
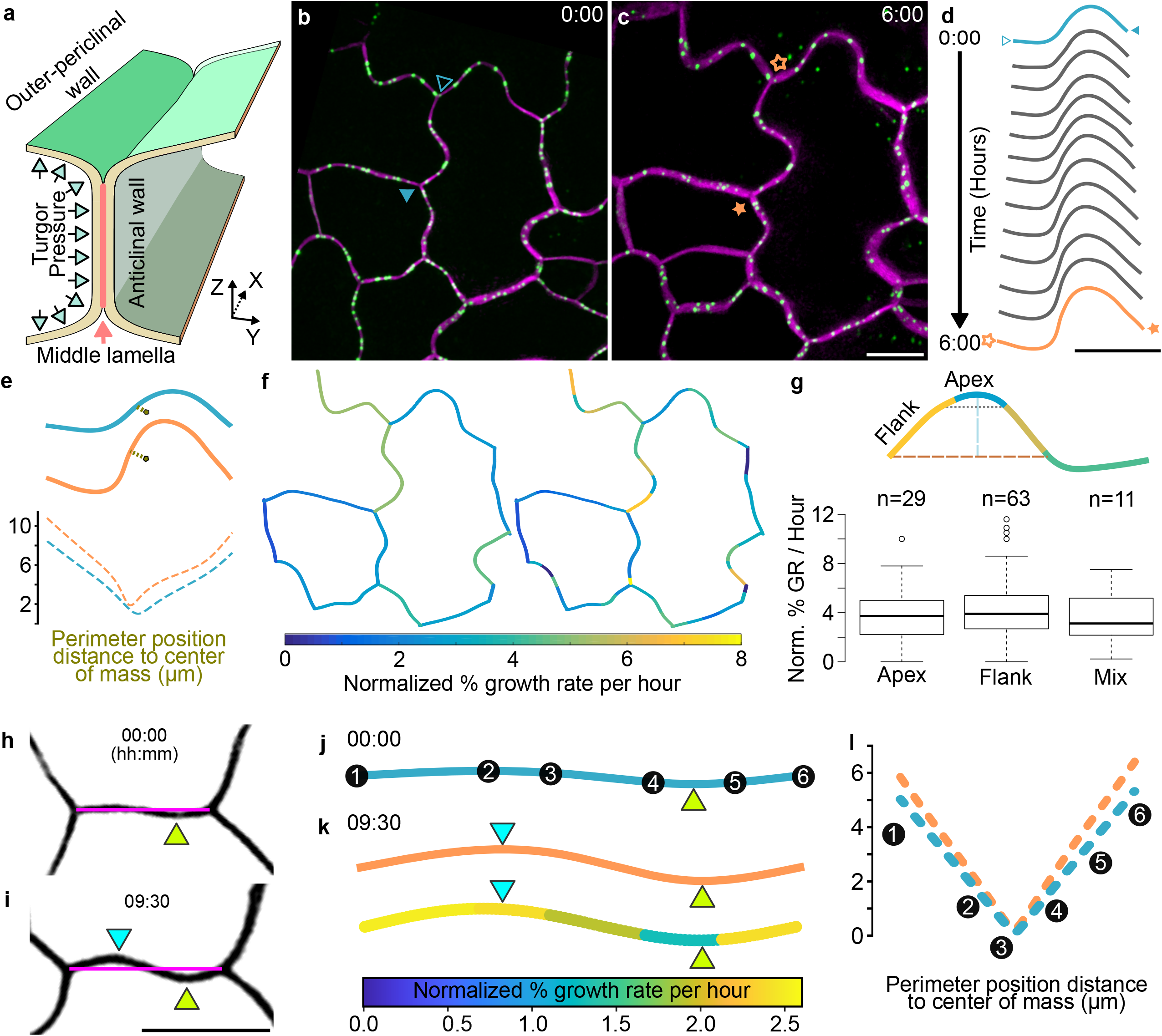
Heterogeneous growth of the cell boundary in developing pavement cells. **a**, Cross-section of cell-cell interface and nomenclature. **c-d**, Example pavement cell field snapshots with tagged plasma-membrane (PIP2:mCherry, magenta) and plasmodesmata fiducial marks (PDLP3:GFP, green), 6 hr interval. **d**, Segment shape change over time from (panel **b** marked segment-green, panel **c** marked segment-magenta). **e**, Segment warping analysis using distance along the segment length to its center of mass after growth (top) which identifies regions of local deformation (bottom). **f**, Spatially mapped growth rates among cell segments in a field (left) and within PDLP-marked subsegments (right). **g**, Lobe geometry does not predict subsegment growth rates. Mean growth rates of a population of subsegments with apex, flank, or mixed localizations. Snapshot of lobing segment 22 min before lobe formation (**h**) and 9:05 hrs after lobe detection (**i**). **j**, Normalized segment length and shape before lobe formation with PDLP positions marked along the segment. **k**, Upper, Normalized segment length and shape at final timepoint showing the previously existing (green arrowhead) and newly formed (cyan arrowhead) lobes. Lower, same as upper but showing normalized subsegment growth rates. **l**, Warping analysis of the cell segment during lobe formation. Perimeter position distance to center of mass plotted for segment shape at lobe detection (**h**,**j**, cyan) and final segment shape (**i,k**, green). PDLPs for initial timepoints are mapped onto the warping analysis plot. Scale bar c, d, i = 10 μm.

Numerous and varied explanations for the mechanism of lobe formation have been proposed^11,13,19,21-24^. The extreme heterogeneity in pavement cell development is one contributing factor. Cells with only subtle undulations and those with a highly convoluted shape coexist in the same leaf sectors for the lifespan of the organ. Further, the growth trajectories of pavement cells cannot be predicted from static images^15,25^, making it impossible to attribute a given pattern of protein localization^12^ or cell wall tensile force^20,22^ to shape change. Historically, cell behaviors were surmised based on static images from a populations of cells^11^. More recent analyses used time-lapse measurements to analyze pavement cell growth^17-19,23,25^. However, lobes originate as a local ~300 nm deflections in the anticlinal wall and appear in the time scale of 10s of minutes^15^, and no parameter or protein that predicts lobe formation has been discovered.

Lobed cells, like others that employ a polarized diffuse growth mechanism, use microtubule-dependent templating of cellulose fibers in the cell wall^26^ to pattern anisotropic expansion^11,13,27^. However, in general, there is a lack of clarity about which subsets of microtubules are morphologically potent. Further, a common assumption is that diffuse growth in plant cells is uniform across the entire cell surface area^28^. However, subcellular growth pattern analyses in several cell types^25,29^ suggest that this might be the exception rather than the rule.

In pavement cells, longstanding models propose that stable microtubules synthesize cellulose arrays or cell wall thickenings that locally restrict growth and cause lobe outgrowths to appear at adjacent subdomains of the cell^11,22^. In Arabidopsis, the microtubule system is unstable and anticlinal microtubules bundling is not associated with local cell wall thickening^15,30^. An alternative growth-promotion model proposes that localized microtubule arrays pattern cellulose microfibrils to generate a patch of local anisotropic expansion that drives symmetry breaking^13^. Even in pavement cells with existing lobes the mechanism by which microtubules promote lobe outgrowth is also unclear. The qualitative enrichment of splayed cortical microtubules at convex cell surfaces has generated different types of growth restriction models^11,19,20,22^. However, a recent quantitative analysis showed that microtubules are only sporadically enriched along subsets of seemingly equivalent convex cell surfaces, and the role of microtubules in lobe outgrowth is unclear^30^.

Additional distinct explanations of interdigitated growth have recently emerged based primarily on the outputs of biomechanical simulations of pavement cells or cell wall components^23,24,31,32^. The finite element (FE) modeling approach is well-suited to analyze plant cell morphogenesis because it simulates stress-strain relationships of thin-walled pressurized shells and can be adapted to any cell geometry^29,33^. Because the material properties and rheology of the cell wall are unknown in most cell types, this approach can make useful predictions. However, the primary liability is that the FE model outputs are highly sensitive to geometry and user-defined variables related to unknown cell wall material properties. Based largely on compositional differences in the opposing anticlinal walls of highly lobed cells Majda et al.^31^ proposed that cell wall stiffness differences across the cell-cell interface drive lobe formation. In another set of simulation studies, a regular patchwork of pectin-controlled cell stiffness differences in the outer periclinal cell wall domains can generate compression and buckling forces in the anticlinal wall and are proposed to cause lobe initiation^23,32^. In another pectin-centric model, vertical bands of pectin in the anticlinal wall^24^, similar to the general pattern of anticlinal microtubule bundles^15^, were identified. The authors created a mechanical model that predicts that the hydration of pectic nanofibers generate forces that cause local cell protrusion. These simulation studies above generated divergent solutions in part because key parameters of the model varied greatly from one model to the next and model predictions were not thoroughly validated with appropriate experimental data. For example, the model of Sapala et al.^22^ was derived and partially validated using large, highly lobed cells that are not in an active phase of lobe initiation^17^.

Here we developed new live-cell imaging and cross-correlation approaches to discover the mechanisms of lobe formation and outgrowth. A multi-channel live cell imaging pipeline was used to accurately graph cell shape and microtubule behaviors at ten-minute intervals for hours highlighting potent transfacial microtubules precede and accurately predict lobe initiation sites. Combinations of mutants, inhibitors, and FE modeling were used to show that the microtubule-cellulose system in an “initiating” cell drives symmetry breaking. Pectin-based adhesion at the cell-cell interface locally couples polarized growth to the adjacent “following” cell to enable interdigitated growth. An FE model based on cell wall properties validated with nanoindentation experiments was used to show that tensile forces in the cell wall provide multi-scale patterning information to localize morphologically powerful microtubules. These results explain how mechanical forces, cytoskeletal, and cell wall systems program polarized growth and provide a general explanation for how plant cells acquire a lobed morphology.

## Results

### Heterogeneous growth and wall tensile force patterns of developing pavement cells

The initial growth analyses focused on the behaviors and properties of the anticlinal wall since it directly reflects shape change, and because growth restriction models predict apical regions of lobes are slow growing^11,19,20,22^. The method was developed using 2 DAG seedlings and fields of early stage, but lobed pavement cells. To directly test for subcellular strain gradients in the anticlinal wall, a two-channel 3D long-term time-lapse imaging pipeline was developed. Cell shape at the anticlinal-periclinal wall interface was accurately quantified with the plasma membrane marker, PIP2mCherry, and local strain gradients were measured using the plasmodesmata-localized protein PDLP3:GFP as fiducial marks (Figure 1a-c; Supplementary Movie 1). By tracking the plasma-membrane coordinates between two 3-way cell wall junctions, it was apparent that lobe height, width, and the length of the adjacent anticlinal wall all increased over time (Figure 1d), a shape change pattern that is consistent with broadly distributed and coordinated cell growth along the lobed cell-cell interface. Segment growth was anisotropic because the boundary changed in shape over the time course (Figure 1e).

Anticlinal cell wall strain was previously assumed to be uniform along the cell perimeter in one previous study^19^. We used PDLP3-marked plasmodesmata as fiducial marks to test for growth rate gradients and correlations with cell shape. Externally applied beads can be used to mark exposed outer periclinal cell surfaces^25^, but it is not possible to get the uniform, high density labeling that is needed for statistical analyses. PDLP3 displacements over time depended on tissue growth (Supplementary Figure 1g-l), and the displacement of paired trackable particles, including the 3-way junctions, were measured at 15 or 30 min intervals for 4.5 to 6 hrs (Supplementary Figure 1a-f). Subsegment growth rates were reliable because in all validated cases the sum of subsegment strains was within 3% of the growth rate that was measured independently using the terminal 3-way junctions. There was a high degree of heterogeneity within individual segments (Figure 1f; Supplementary Figure 2j). To determine if there were differences in the growth rates of lobe apex or flanks, segments were assigned to lobe sub-domains based on standardized landmarks (Figure 1g – top). A population-level analysis of the apex, flank, and mix sub-segments showed no significant difference in anticlinal wall growth rates in these subdomains (Figure 1g – Bottom).

With this imaging pipeline it was not technically feasible to conduct similar statistical analyses of lobing segments because PDLPs do not provide the spatial resolution needed to analyze the subdomains of emerging lobes and it was difficult to get a large number of time-lapsed datasets from lobing segments. Nonetheless we were able to analyze 3 lobing segments and determine if their growth behaviors were similar to those of lobed cells. Lobing segments displayed growth heterogeneities that were similar to those of lobed segments (Figure 1h-k and Supplementary Figure 1m-o). Interestingly, growth rates in the regions of newly formed lobes varied greatly between segments residing in domains with the highest (Figure 1k) and lowest (Supplementary Figure 1n) growth rates. Warping patterns of lobing segments were variable, but not dominated by the locales of lobe initiation (Figure 1l; Supplementary Figure 1m-o). These data indicate that polarized growth that generates new lobe features is superimposed upon asymmetric growth patterns that occur at the cell segment scale.

To test for a correlation between growth rates and tensile stress, a 3D finite element (FE) model of the cell clusters was generated using cell geometries obtained directly from the live-cell image data on the cells of interest (Figure 2a,b; Supplementary Figure 2j,k). The FE model simulates the cell as a thin-walled pressurized shell with wall properties based on an isotropic, neo-Hookean hyperelastic, material behavior (see Methods) (W. Li, J.A Turner, Biorx ID to be provided). Based on direct measurements from TEM images, the model includes cell wall thickness of 300 nm for the outer periclinal wall and 40-50 nm for each of the paired anticlinal walls, and uses experimentally validated turgor pressure measurements and realistic cell wall modulus values on the order of a few hundred MPa. The model also includes an adherent layer to simulate the middle lamella and independent anticlinal walls in each cell that respond to tensile forces (Figure 2b). The unpaired outer periclinal wall fully bears turgor pressure and the in-plane tensile forces pull upward on the anticlinal wall, and the model generates quantitative predictions of cell wall stress magnitude and direction as a function of location in the cell.

**Figure 2.**
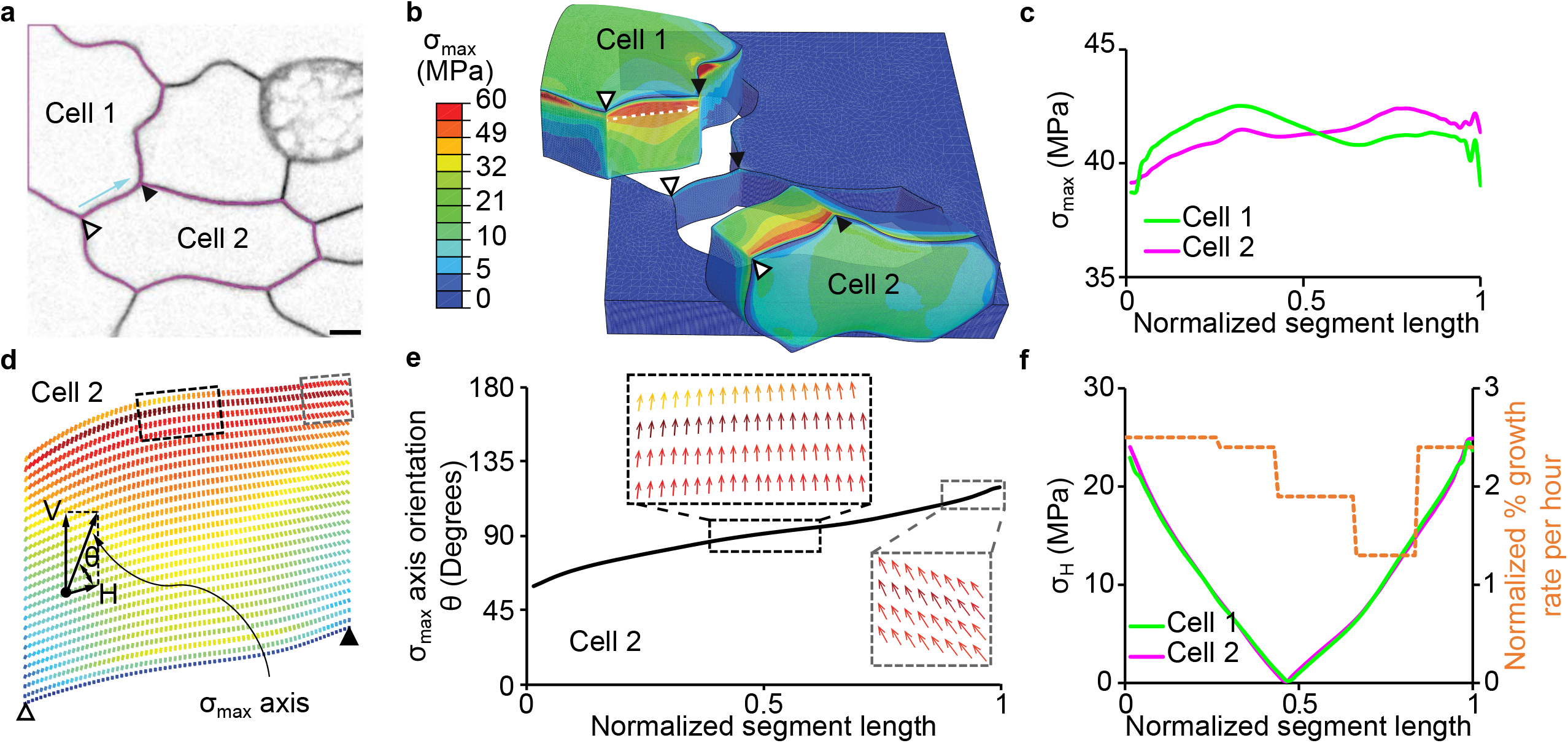
**a**, Cell field used to create the FE model of a pavement cell cluster; arrow, segment used to compare strain and wall tensile stress patterns. **b**, FE model of two cells showing the variation of maximum principal stress. **c**, Distribution of maximum principal stress (Max PS) in the anticlinal walls. **d**, Orthographic projection of a representative curved lobing segment. The orientation of each short line represents the Max PS axis which can be divided into its horizontal and vertical components. **e**, Max PS orientation across the segment. Dotted box show the normal view of the anticlinal wall with arrows reflecting the magnitude and direction of wall stress and orientation. **f**, Correlation analysis of segment strain and horizontal component of anticlinal wall Max PS plotted graphically as a function of cell position. Scale bar = 5μm.

Because of their large size, many cell pairs of interest were not completely captured in the image field. The technical issue of effects of partial cell volumes on stress distributions was addressed in a series of sensitivity analyses (Supplementary Figure 2a-e) that showed anticlinal wall stresses based on partial cells were largely unchanged (< 10 %) except when the majority of one cell was missing. Most critical was the area and location of the maximum height of the periclinal wall, which could be estimated from image data for the pairs included in this work. An FE model of the lobing cell pair in Figure 1h-l was generated based on estimated area for the partial cell 1 using the image data and the maximum height profile of the real cell (Supplementary Figure 2f-i). The estimated area of cell 1 of 692 μm^2^ is similar to the mean cell area of 634 μm^2^ in 1 DAG cotyledons (Supplementary Table 1). The maximum principle stress (σ_max_) profiles for the two opposing anticlinal walls at the lobing interface were extracted from the FE model (Figure 2b). Along cell segment length there was a general trend for the magnitude of σ_max_ to decrease approaching 3-way junctions (Figure 2b,c; Supplementary Figure 2k-o). The mean σ_max_ values along a segment decreased when averaged over increasing depths because of the drop in stress away from the periclinal wall, and the shapes of the profiles were most consistent near the top of the wall (Supplementary Figure 2e). Because our subsequent analyses focus on the interface of the outer periclinal and anticlinal walls, we report σ_max_ profiles as the average values from the upper 1/8th of the anticlinal wall.

As expected, the σ_max_ tended to be oriented perpendicular to the outer periclinal wall; however, off axis tensors were frequently observed near 3WJ and in some regions of high cell curvature (Figure 2d-e; Supplementary Figure 2l-o). The vertical component of σ_max_ is not the primary driver of anticlinal wall expansion since cell growth occurs primarily in the plane of the leaf, and this may reflect the vertical orientation of microtubules^15^ and the expected cellulose-dependent anisotropy of the anticlinal wall. As σ_max_ deviated increasingly from vertical, the horizontal component (σ_H_) of this tensor increased, and displayed a high degree of spatial variability (Figure 2d-f; Supplementary Figure 2l-o). Interestingly, within each of the 5 FE model segments for which σ_H_ profiles were generated, the σ_H_ values correlated with the sub-segment growth rates measured using PDLP3 (Figure 2f, Supplementary Figure 2l-o). This suggests that the direction of σ_max_ is a key determinant of local growth rates, and that within a segment the anticlinal wall is highly anisotropic and has similar material properties along its length. There may be variability in wall material properties between segments as the growth rates between segments differed greatly even though simulated σ_H_ values were similar. These modeling and growth kinematics depict a general planar cell expansion mechanism in which σ_H_ defines anticlinal wall growth rates, and material properties are maintained via coordinated cell expansion and new wall synthesis. Lobe initiation occurs within this cellular-scale biomechanical context, and the next major challenge was to determine what gene activities and molecular functions control lobe formation.

### The central importance of the microtubule-cellulose system during lobe initiation

An initial model for lobe initiation was based on subcellular gradients of auxin and trans-cellular ROP small GTPase signaling^21^. However, recent genetic and localization studies failed to detect a pavement cell function for the major components of this auxin-based pathway^14,15^. Given the divergent models proposed for lobe formation and the recently proposed central roles of pectin^23,24^, we screened a wide collection of morphology mutants and inhibitors for lobe number defects. Fully expanded wild-type pavement cells have a low area to perimeter ratio with a mean circularity value of 0.26 and a mean lobe number per cell of 16 (Table 1; Supplementary Figure 3a). The *anisotropic1 (any1)* mutation is in a *CELLULOSE SYNTHASE1* complex subunit and causes defects in microfibril organization and polarized growth^34^. *Any1* had a much simpler overall shape and a reduced lobe number both on a per-cell and cell area normalized basis (Table 1; Supplementary Figure 3f). KATANIN1 is a known microtubule severing protein and has been proposed to link cell wall tensile force with morphogenesis^35^. The *ktn1* mutant had a simpler shape compared to the wild type, but no detectable reduction in lobe number. Similar results were obtained with the microtubule plus-end binding mutant *clasp* (Table 1; Supplementary Figure 3b,c). The lack of a lobe initiation phenotype for *clasp* might be due to the radius of curvature of 2.59 ± 0.27 μm (8 cells) at the transition from the outer periclinal wall to the anticlinal wall in 2 DAG pavement cells. This exceeds the radius of curvature threshold of 2.5 μm, below which CLASP-dependent stabilization of transfacial microtubules operates^36^. These phenotypes could be explained by a defect in the maintenance of lobe outgrowth rather than lobe initiation, similar to what has been shown for the microtubule-binding protein BPP^30^. A possible effect of the microtubule plus-end binding MOR1 on pavement cell shape has been reported^37^. In our assay, the temperature-sensitive *mor1-1* allele had a reduced lobe number when it was shifted to the non-permissive temperature after germination (Table 1; Supplementary Figure 3h,j). These data are consistent with the proposed importance of microtubules during lobe initiation, but MOR1 is not a useful molecular marker for the specific microtubules that control lobe initiation because its mutation causes global and severe defects in the cortical microtubule network^38^. Pectin has been implicated as a patterning molecule in lobe initiation^23,24^. The *quasimodo2 (qua2)* mutant has reduced pectin synthesis^39^. The *qua2* mutant had a reduced cell size and number of lobes per cell, but an increased lobe number when normalized to the cell area. There was no indication that *qua2* reduced lobe number when cell protrusions at 3-way junctions were manually removed to specifically quantify type I lobes^40^; the mean lobe number/area x 1000 was 1.71 ± 0.76 for *qua2-1* and 1.13 ± 0.28 for similarly sized wild-type cells. Surprisingly, mutation of exocyst complex subunit *exo84b*, which is involved in targeted secretion, increased lobe number, pointing to additional pathways that limit lobe initiation (Table 1; Supplementary Figure 3g).

**Table 1.**
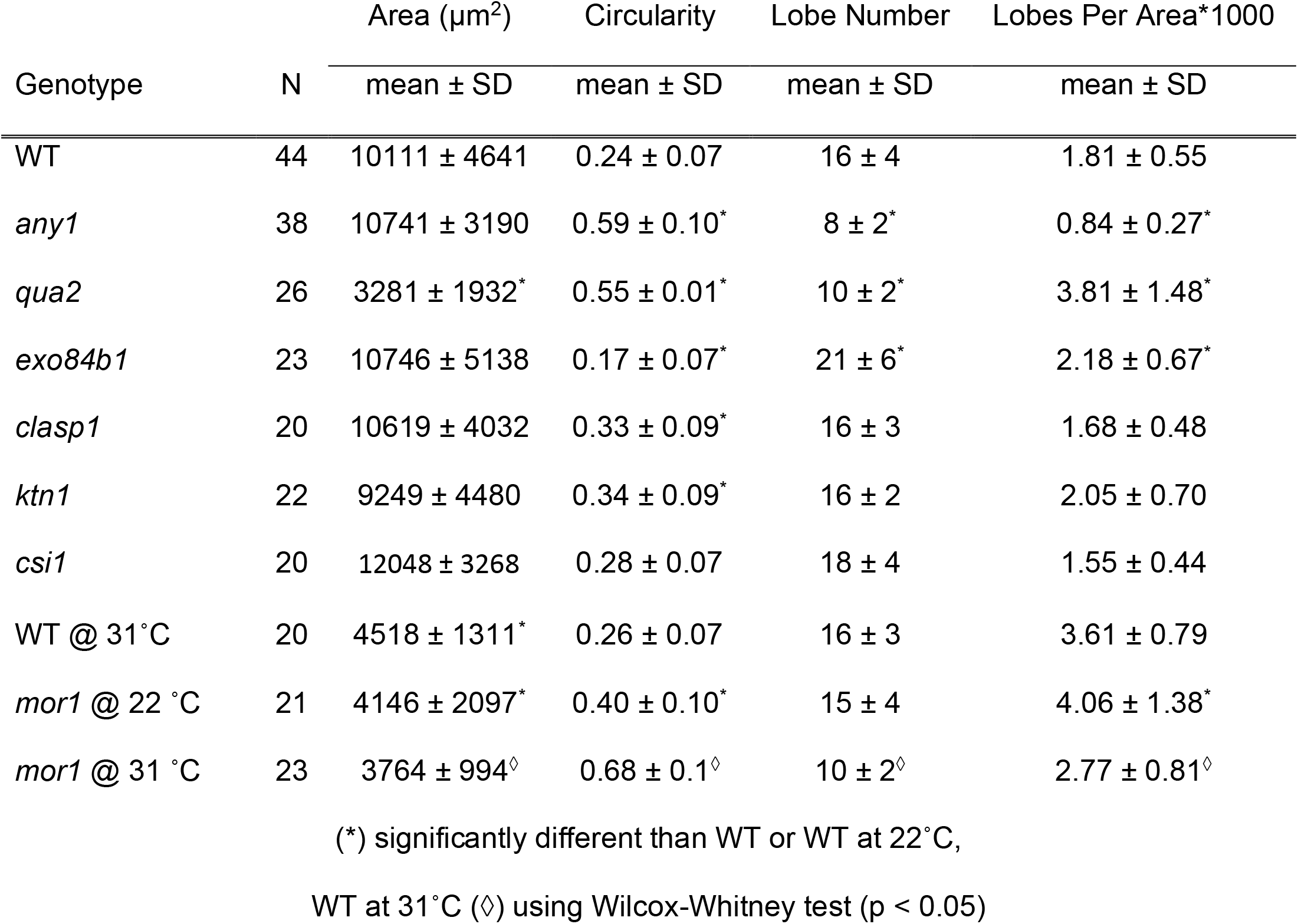
Mutations that affect microtubule polymerization and cellulose synthesis reduce pavement cell lobing

| Genotype | N | Area ($\mu\text{m}^2$) | Circularity | Lobe Number | Lobes Per Area*1000 |
| --- | --- | --- | --- | --- | --- |
| | | mean ± SD | mean ± SD | mean ± SD | mean ± SD |
| WT | 44 | 10111 ± 4641 | 0.24 ± 0.07 | 16 ± 4 | 1.81 ± 0.55 |
| *any1* | 38 | 10741 ± 3190 | 0.59 ± 0.10* | 8 ± 2* | 0.84 ± 0.27* |
| *qua2* | 26 | 3281 ± 1932* | 0.55 ± 0.01* | 10 ± 2* | 3.81 ± 1.48* |
| *exo84b1* | 23 | 10746 ± 5138 | 0.17 ± 0.07* | 21 ± 6* | 2.18 ± 0.67* |
| *clasp1* | 20 | 10619 ± 4032 | 0.33 ± 0.09* | 16 ± 3 | 1.68 ± 0.48 |
| *ktn1* | 22 | 9249 ± 4480 | 0.34 ± 0.09* | 16 ± 2 | 2.05 ± 0.70 |
| *csi1* | 20 | 12048 ± 3268 | 0.28 ± 0.07 | 18 ± 4 | 1.55 ± 0.44 |
| WT @ 31°C | 20 | 4518 ± 1311* | 0.26 ± 0.07 | 16 ± 3 | 3.61 ± 0.79 |
| *mor1* @ 22 °C | 21 | 4146 ± 2097* | 0.40 ± 0.10* | 15 ± 4 | 4.06 ± 1.38* |
| *mor1* @ 31 °C | 23 | 3764 ± 994$\diamond$ | 0.68 ± 0.1$\diamond$ | 10 ± 2$\diamond$ | 2.77 ± 0.81$\diamond$ |

(\*) significantly different than WT or WT at 22°C,
WT at 31°C ( $\diamond$ ) using Wilcox-Whitney test (p < 0.05)

We conducted a series of inhibitor experiments to analyze microtubule, cellulose, and pectin systems in the context of lobe initiation (Supplementary Figure 3k-q; Supplementary Table 1). In the interval from 1-to 2-days after germination (DAG), wild-type cells generated ~5 lobes/cell in the presence or absence of 0.1% DMSO (a solvent control for oryzalin). Treatment with oryzalin, a microtubule-depolymerizing drug, and isoxaben, an inhibitor of CesA, decreased lobe initiation rates significantly but not the cell areal growth rates relative to 2 DAG controls. Treatment with 0.2% pectinase, which selectively degrades homogalacturonan, had no detectable effect on growth or lobe formation (Supplementary Table 1). We show later in this study that this concentration of pectin had a strong effect on cell-cell adhesion. These results further reinforce the primary importance of the microtubule and cellulose systems for lobe initiation. The lobing defect was due to microtubule-biased directionality of CESA trajectories in the plasma-membrane^41^ because time-lapse imaging of YFP:CesA6 and mCherry:TuA5 showed that many linear CESA tracks mirrored the spatial distribution of microtubules (Supplementary Figure 3r-t). High spatial and temporal time-lapses of CesA and microtubules at the anticlinal wall are not possible due to photobleaching. Resliced snapshots showing the face-view of the anticlinal wall show evidence that the microtubule-CesA colocalization is consistent at this location (Supplementary Figure 3u - right panels). The *csi1* mutant was tested for lobe defects based on its known involvement in coupling CESA to cortical microtubules^42^. Surprisingly, there was no phenotypic difference between the *csi1* mutant and wild-type populations (Table 1, Supplementary Figure 3e). This result reflects the fact that colocalization of CESA and microtubules was retained at both the outer-periclinal and anticlinal walls in *csi1* (Supplementary Figure 3v-y), and likely involves additional pathways or CSI1-like genes. Collectively these data suggest that microtubule-dependent patterning of cellulose fibers in the wall is an essential activity to pattern interdigitated growth.

### Transfacial microtubules predict the location and direction of lobe formation

High-resolution imaging of the plasma-membrane and microtubules consistently detected perpendicular anticlinal microtubule bundles that splayed along the cortex of the outer- and inner-periclinal cell surfaces (Figure 3a,b). This suggested that microtubules function symmetrically at both periclinal cell surfaces. Time-lapsed analyses of anticlinal wall tilt showed that the anticlinal wall remained straight during lobe formation and outgrowth (Supplementary Figure 3z-ee) and reflected symmetrical growth of the outer and inner periclinal walls.

**Figure 3.**
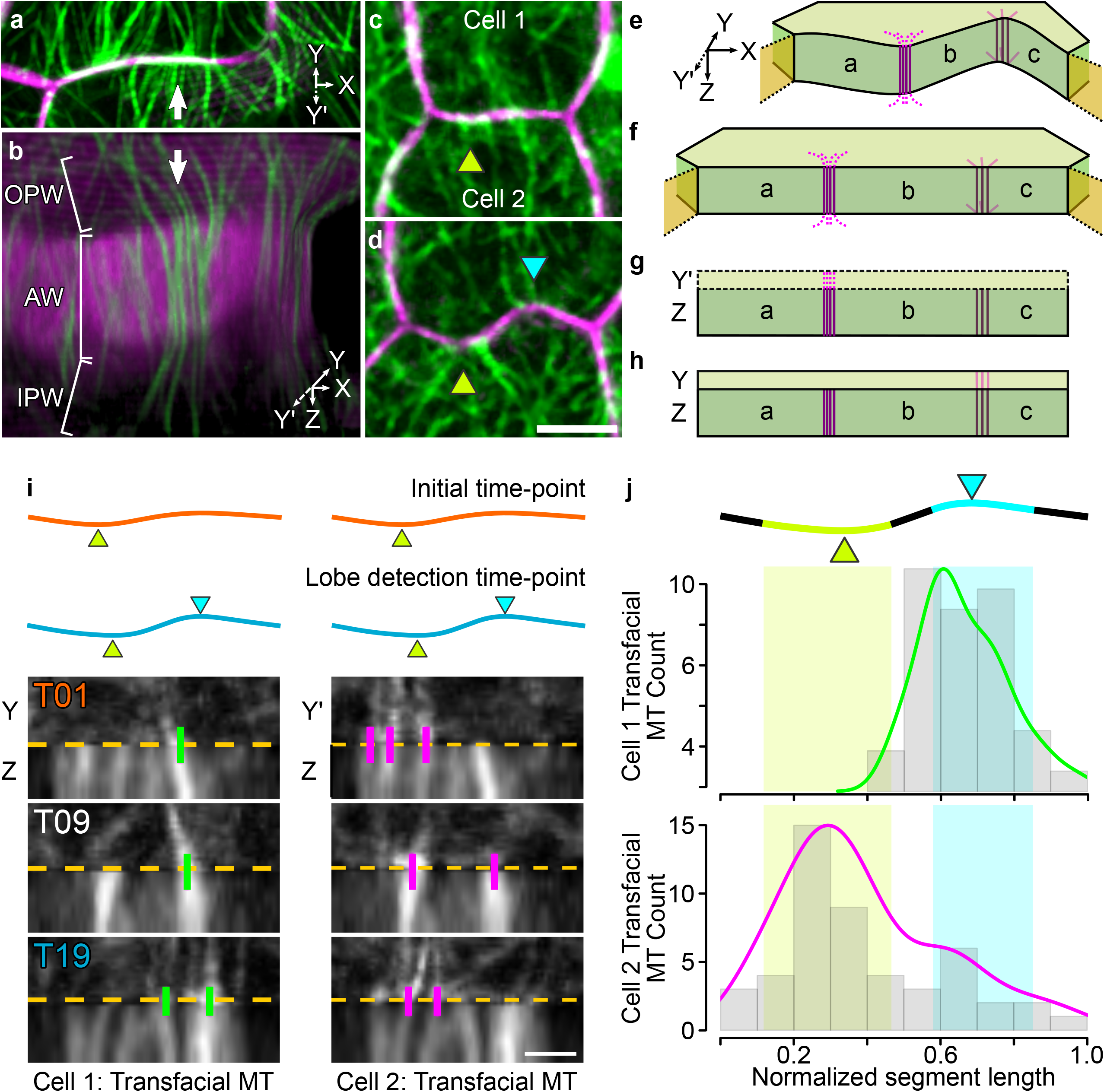
Transfacial microtubules predict lobe initiation sites. **a**, Top-view of a segment that will form a lobe; plasma-membrane (magenta) and microtubules (green) rendered in 3D. **b**, where microtubules (MT) are seen splayed from the anticlinal wall to the inner- and outer-periclinal walls. **c-d**, Initial and final snapshots of a segment that contains an existing (green arrowhead) and a newly formed lobe (cyan arrowhead). **e-h**, Method for transfacial microtubule analysis (see methods for details) resulting in an image montage of the anticlinal and outer-periclinal walls for the adherent cells (**g, h**); a,b,c: subregions of the coupled anticlinal walls; Y’: periclinal wall of cell 2; Y: periclinal wall of cell 1. **i**, Transfacial microtubules at a lobing interface in cell 1 (left) and cell 2 (right). Lobing segment shape (top) at the initial (orange) and lobe detection (cyan) timepoints. Montaged images of anticlinal and outer-periclinal walls at three indicated time points with transfacial microtubules indicated with vertical bars (green - cell 1, magenta - cell 2). **j**, Histograms of transfacial microtubules location before lobe detection showing enrichment in cell 1 near the lobing region (cyan) and cell 2 (yellow) near the region of an existing lobe. Scale bar = 5 μm for **c-d**; 2 μm for I.

To determine how microtubules control lobe formation, we took a multivariate long-term live-cell imaging approach using cells in which microtubules and the plasma membrane were marked (Supplementary Movie 2). Prior analyses of microtubules and lobing segments were based on either snapshots of already lobed cells^11,12^ or time-lapsed analyses at time intervals of 1 hr or greater^15^ to 6-hr^19,23^, which is not sufficient because lobe initiation occurred at time scales of tens of minutes (Figure 3c,d; Supplementary Movie 3). Correlation analyses of time-series microtubule imaging consistently showed that the microtubule network turned over in approximately 10 min (Supplementary Figure 4a-f). Because 20-25 planes were needed to image the outer periclinal and most of the anticlinal cortex of lobing cells, bleaching was a major challenge. Therefore, we selected 10 min intervals that allowed us to image over ~4-to 8-hr intervals at low laser power and capture most of the detectable differences in the microtubule network, and at this sampling frequency, 32 % of all transfacial microtubules were detected two or more consecutive time-points. In this paper, the term microtubule refers to either single microtubules or microtubule bundles.

In order to graph outer periclinal and anticlinal microtubule behaviors as a function of the location and timing of lobe detection, candidate cell segments were tested for lobe formation events post-acquisition using the prominence method^15^. Opposing patches of the outer periclinal cell cortex at the future site of lobe detection do not differ in microtubule density^15^. We found that the orientation and coherency of microtubules in the opposing outer periclinal cell patches also varied wildly as a function of time prior to lobe initiation (Supplementary Figure 4g-n). This is not unexpected because many periclinal microtubules are randomly oriented and enter and exit the region of interest from any direction. Despite the high variability of the microtubule network, summed projections of the time series data consistently revealed hot spots of microtubule signal at or near the future site of lobe initiation (Figure 4b, Supplementary Figure 5p,t).

**Figure 4.**
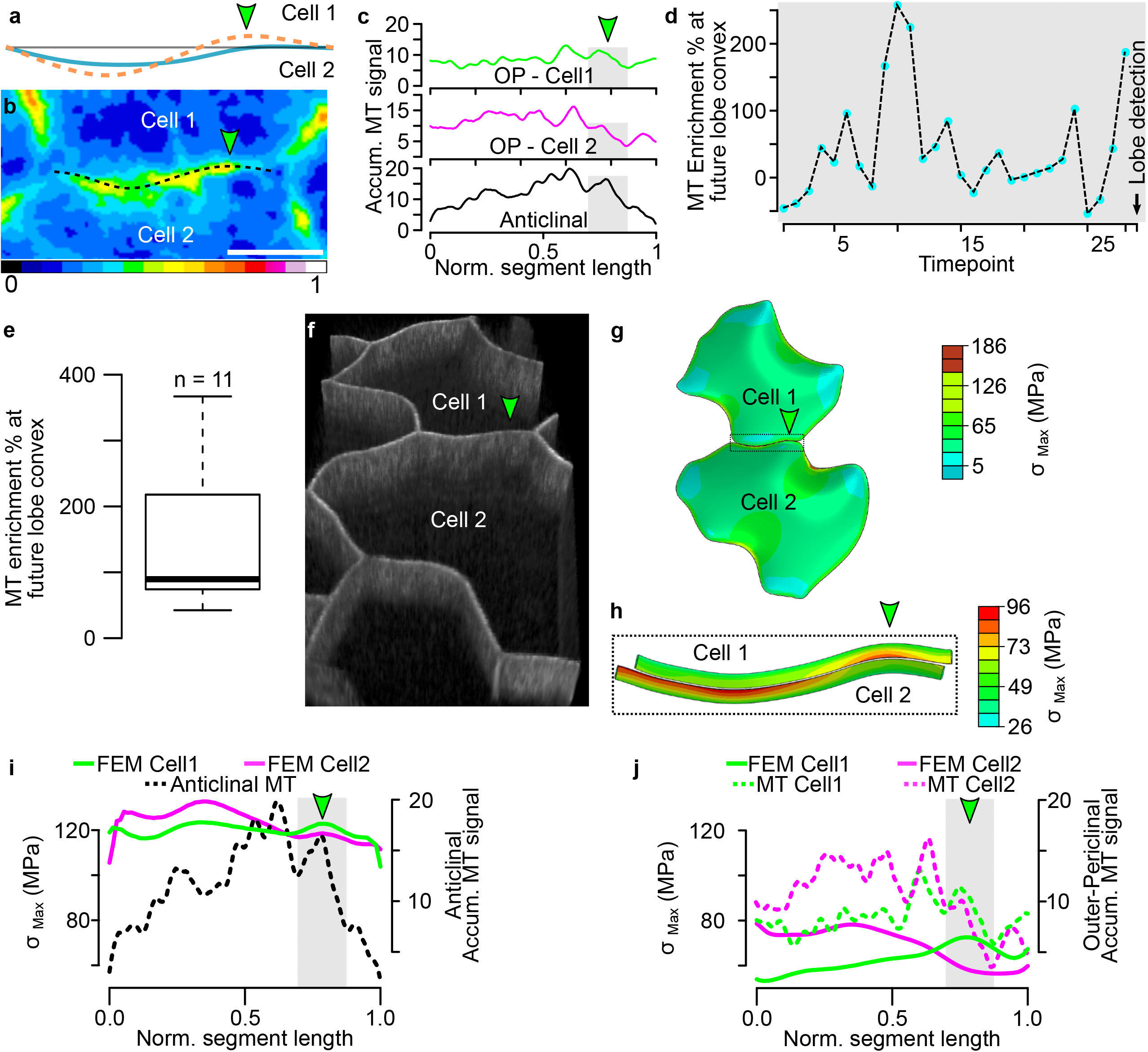
Microtubule signals are concentrated at regions of elevated cell wall tensile stress. **a**, Segment shape at the start of the time-lapse (blue) and at the time point of lobe detection (dashed - orange) that is used for the analysis on (**c-d, f-j**). Green arrowhead denotes location of new lobe. **b**, Heatmap of normalized microtubule signal intensity before lobe initiation shows a hotspot at the future lobe initiation site (green arrowhead). **c**, Accumulated microtubule signal intensity at the periclinal cortex of cells 1 (green line), cell 2 (magenta line), and the anticlinal cortex (black line) before lobe is detected. Future lobe site region shaded grey. **d**, Plot Microtubule (MT) enrichment at the future convex side of the new lobe over time. 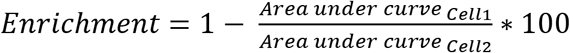. **e**, Boxplot of microtubule enrichment at future lobe convex before lobe detection, n is the number of lobing segments analyzed. **f**, 3D rendering of the cell pair used to construct the FE model shown in (**g**) before the new lobe was detected (green arrowhead is future lobe region). **h**, Zoom-in view of the segment from (g - dotted rectangle) with maximum principal stress gradients mapped on the outer periclinal walls adjacent to the anticlinal walls. **i**, The accumulated anticlinal microtubule signals before lobe detection (dashed line) and anticlinal wall stress distributions from cell 1 (green) and cell 2 (magenta) from FEM are plotted as a function of normalized segment length. **j**, The accumulated periclinal microtubule signals (dashed lines) and periclinal wall stress distributions (solid lines) for cell 1 (green) and cell 2 (magenta) are plotted as a function of normalized segment length. Scale bar = 5μm.

Transfacial microtubules that span the anticlinal and periclinal walls are strong candidates to control lobe initiation because they can locally define cell wall properties across both cell faces^13^. To quantify transfacial microtubules, 3D images of the lobing interface were straightened, and the outer periclinal walls of the future-convex and future-concave cells were rotated and aligned with resliced images of the anticlinal surfaces to create a 2D projection of the two faces of each cell (Figure 3g,h; Supplementary Movie 4). Transfacial microtubules were defined as those that had continuity between the outer periclinal and anticlinal cortex (Figure 3i - magenta and green tags). The location and frequency of transfacial microtubules in both cells were plotted prior to lobe detection. Segment lengths were normalized between time points to account for growth during the time course (Figure 3j). The future convex cortical domain was defined as a subsegment equal to the width of the lobe feature at its ½ maximal height centered at the lobe initiation site^15^. Notably, there was an obvious enrichment of transfacials in the future convex cell, with the peak centered near the location where the new lobe will emerge (Figure 3j). Supplementary Figure 4o-s summarizes identical manual quantification data for 5 additional independent lobing events with similar behaviors.

To increase the throughput of our microtubule-cell shape cross-correlation analyses a semi-automated approach was developed. Anticlinal microtubules are perpendicular to the leaf surface, and their localization can be quantified by summing pixel intensities along vertical sectors of the image^15^. Because the dominant orientation of transfacial microtubules in the outer periclinal cell surfaces in our processed images are also parallel and orthogonal to the anticlinal wall (Figure 3i; Supplementary Figure 5c,f), a line profile of the summed microtubule intensity in a vertical sector was used to graph microtubule signals on this cell face as well. Microtubule signals were normalized from 0 to 1 and plotted cumulatively over the time course up to the time point prior to lobe initiation (Supplementary Figure 5g-n). The plots were noisy compared to the manually scored images because all microtubule signals along the segment were included; however, there was a consistently elevated microtubule signal in the future convex periclinal domain compared to an equivalent region at the future concave domain (Figure 4c,e; Supplementary Figure 5n,s,w). In all datasets, there was a large degree of temporal variability in the degree of microtubule enrichment, with the highest values often occurring in a few sequential time points. Similar trends in temporal variability were observed in manually scored time-series data (Supplementary Figure 4t-y).

The mean degree of microtubule enrichment prior to lobe initiation varied greatly among future convex domains (Figure 4e; Supplementary Figure 5o), indicating that there is no single microtubule behavior that is sufficient to predict lobe formation events. However, this microtubule organization has the power to influence cell shape. In one time series, the symmetry-breaking event operated against a clearly existing broad concave bulge (Supplementary Figure 5x-aa). This asymmetric and localized lobe initiation event cannot be explained by a buckling event as previously claimed^23^. These analyses demonstrate that transfacial microtubules persistently accumulate at a cortical site that accurately predicts the location and direction of lobe initiation.

### Cell wall tensile forces predict locations of microtubule enrichment

The mechanism that determines the spatial pattern of cellular interdigitation is not known. Lobe formation conforms to a minimum spacing rule^40,43^, and we qualitatively observed in our time-lapse analyses that lobes never form within an existing lobe and tend to appear at the midpoint of straight regions of the anticlinal wall that are greater than 6 μm. Biomechanically it is possible that this geometric bias reflects local regions of elevated cell wall tensile force. Prior FE modeling and microtubule imaging suggest that microtubule orientation is correlated with cell wall stress^20^, and our FE simulations of a lobing segment indicated that σ_max_ profiles can differ between adjacent anticlinal cell walls (Figure 2c).

To determine if patterns of cell wall tensile forces correlated with microtubule behaviors that relate to lobe initiation, a 3D model was created based on image data of lobing cells in the same manner as in Figure 1 (Figure 4f-h). As expected in lobing segments^15^, cortical microtubules along the anticlinal walls were highly oriented parallel to σ_max_ tensors (Supplementary Figure 5a-f). The spatial distributions of σ_max_ values were quantified in the upper 1/8 of the anticlinal (Figure 4i) and outer periclinal (Figure 4j) walls adjacent to the cell-cell junction. The stresses along the anticlinal wall were around 120 MPa, and cell 1 had a small peak of anticlinal wall stress centered at the future convex region of the cell (Figure 4i). The sectors of the two adjacent periclinal walls along the cell segment had a more offset pattern of stress. In cell 2, the stress maximum correlated with the convex curvature of an existing lobe (Figure 4j). Cell 1 had a peak of slightly elevated stress that was centered on the future lobe location. FE models were constructed for 2 additional lobing segments, and the same pattern held. Stress profiles were similar along the anticlinal wall, and the future convex cell had a locally elevated tensile force in the periclinal wall that was at or near the future convex cortical domain and the site for microtubule enrichment (Supplementary Figure 5r,s,v,w). The stress predictions of the model are approximations of the real cell behavior because cell wall properties were assumed to be homogeneous, but these analyses were consistent with tensile force acting as an upstream patterning element during interdigitated growth. In this scenario, microtubules convert tissue- and cellular scale tensile force patterns into a polarized growth process that leads to lobe formation.

### A cell-autonomous system to analyze wall stress, microtubules, and lobing

We next used cell-cell adhesion mutants and pectinase treatment to decrease cell-cell adhesion and enable cell autonomous growth to generate cells with altered shapes and tensile stress distributions. If the future convex cell uses the microtubule-cellulose systems to initiate lobe formation, then a central function of pectin and the middle lamella could be to mechanically couple the initiating cell with complimentary polarized growth in the “following” cell. We had noticed previously that cell-cell adhesion mutants *qua2* had unusual invaginations at the cell periphery (Figure 5a). We wanted to examine the ultrastructure of furrows using electron microscopy, but *qua2* seedlings were too fragile to survive the sample processing protocol. As an alternative, we turned to the known cell-cell adhesion mutant *dis2*^44^, which also occasionally displayed clear invaginations at the cell periphery. We developed a short-term pectinase treatment protocol that greatly increased the rate of furrow formation in *dis2* (Figure 5b,c) and the wild-type (Figure 5d). Invaginations were not observed in wild-type control samples that were not pectinase-treated. In medial longitudinal thin sections through *dis2* furrows closely opposed cell walls from the same cell and a widened bulb at the apex was observed (Figure 5e). The apex was frequently enriched with cortical microtubules that could be seen in the cross-section. This cell geometry, microtubule organization, and ultrastructure exactly matched the invaginations that have been detected in lobed mesophyll cells^45^. Furthermore, a previous analysis of cell-cell adhesion in the context of organ development included a time-lapse image series documenting the slow inward progression of a furrow-like structure^46^. Furrows are generated over time scales of hours and reflect a shape deformation that is generated by slow irreversible growth.

**Figure 5.**
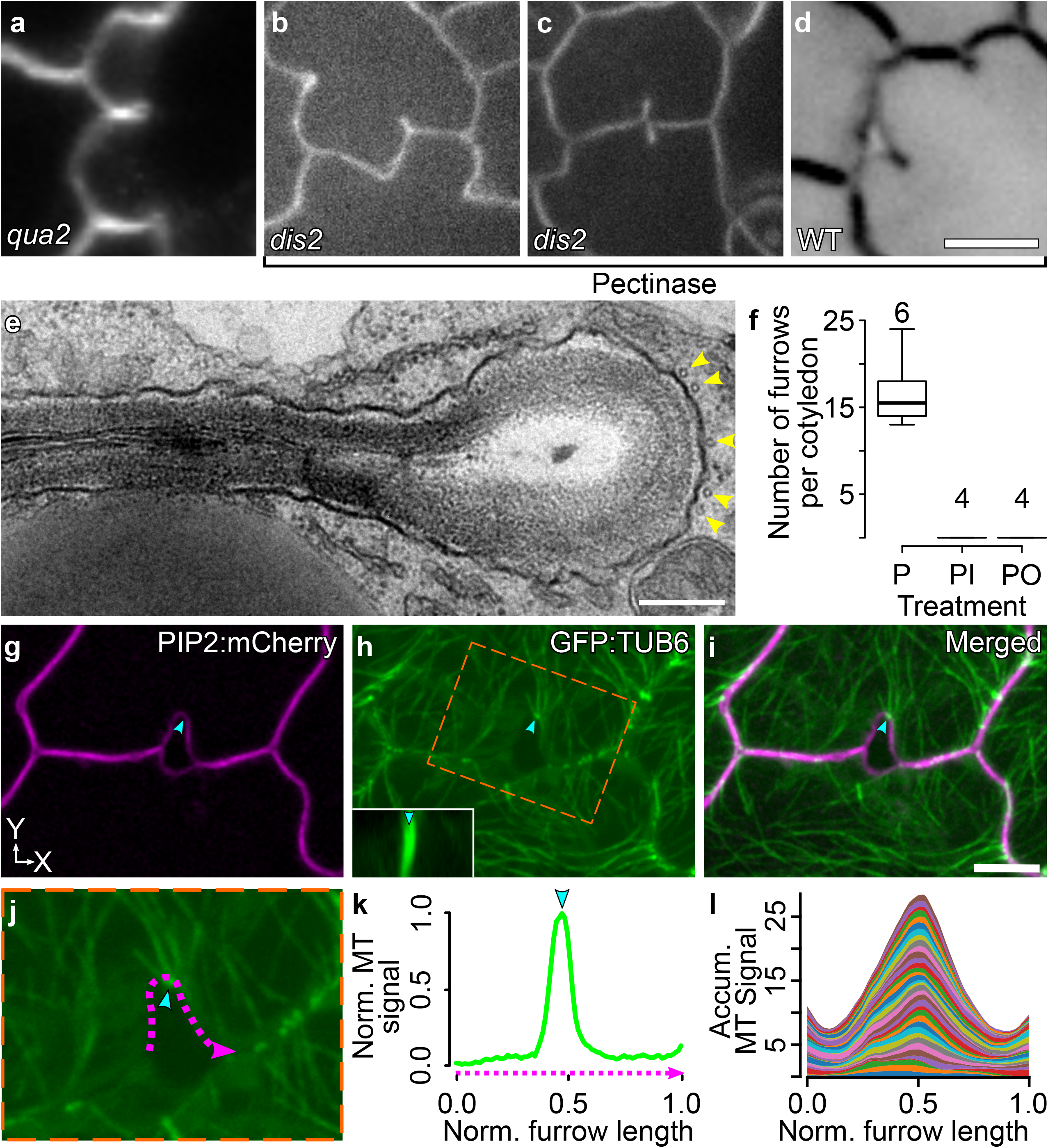
A system for inducible cell autonomous lobing. **a**, The pectin-deficient mutant, *qua2-1*, develops cell invaginations in pavement cells. **b-c**, Pavement cell furrows in the cell-cell adhesion mutant *dis2*. **d**, Furrow formation induced in wild-type by 0.2% pectinase treatment. **e**, TEM analysis of an induced furrow in *dis2,* arrowheads, cortical microtubules. **f**, Box plots of the number of furrows/cotyledon of wild-type, pectinase-treated seedlings. P - pectinase, O - Oryzalin, and I - Isoxaben at 0.2%, 1 nM, and 5 μM concentration respectively. The number above boxplots is the number of cotyledons analyzed. **g**, Face on view of a nascent furrow developing from pectinase-induced delamination. (**h-i**) Splayed microtubules at the apex of a nascent furrow. **j**, Inset region of (**h**), furrow outline (magenta), and microtubule signal maximum (cyan arrowhead). **k**, Plot of normalized furrow position and microtubule (MT) signal to enable a population-level analysis. **l**, Population-level analysis of microtubule signals and location along nascent furrows (n = 36). Panels **g-l** were generated using wild-type pectinase-treated cells. Scale bars = 10 μm for **a-d**; 250 nm for **e**; 5 μm for **g-i**.

The progressive invaginations of pectinase-induced furrows were generated by microtubule- and cellulose-dependent processes because the delaminations and furrows were greatly reduced by oryzalin and isoxaben at concentrations that did not inhibit cell expansion (Figure 5f). The similar growth patterns and inhibitor sensitivities of furrows and lobe formation during normal cotyledon development (Supplementary Table 1) indicate that these two morphogenetic processes were analogous and employed similar cell shape control mechanisms. Interestingly, furrows usually were unidirectional, but bidirectional furrows were occasionally observed (Figure 5c), indicating that at least in some instances the directionality of the invagination is not predetermined by existing cell wall material properties at the time of pectinase treatment.

To analyze microtubule localization in furrows more quantitatively we identified the rare early-stage invaginations in which an asymmetrically expanded feature was present at the site of pectinase-induced delamination. In nascent furrows the delamination was not symmetrical, and one cell boundary was more invaginated compared to its neighbor, the opposing cell walls were not appressed, and there was no terminal bulb (Figure 5g). Similar to convex sites of lobing pavement cells the curved apex of the nascent furrows frequently had anticlinal microtubule bundles that splayed out at the anticlinal/periclinal wall interface (Figure 5h,i). When line scans of microtubule signal intensity were plotted as a function of position along the plasma membrane boundary for a population of nascent furrows there was a strong bias for microtubule location at the apex (Figure 5j-l). This bias in microtubule localization in the more polarized cell at delamination likely reflects the microtubule-dependent generation of the asymmetric cell shape, that perhaps includes changes in tensile force that could provide positive feedback over the extended timescales of cell invagination (see below). This cell autonomous lobe/furrow formation system demonstrates the existence of an initiating cell that uses the microtubule and cellulose systems to drive cell ingrowth and the middle lamella couples that asymmetric growth to the adjacent cell.

### Cell wall tensile force patterns define the location and pattern of furrow formation

If furrow formation parallels interdigitated growth then cell wall tensile stress patterns should influence their patterning and development. In this instance, pectinase treatment compromised cell-cell adhesion and provided a highly useful experimental manipulation that enabled cells to respond differently to endogenous cell wall stress patterns and generate unique cell shapes with altered stress distributions. FE modeling was used to analyze both of these processes by constructing a model based on the image data of a cell pair with a nascent furrow (Figure 6a). We found furrow location was strongly biased to the mid-region of the segments (Figure 6f), similar to initiating lobes. In the furrowing cell pair, the location and directionality of furrow corresponded to the stress maximum of the future convex cell when the FE model boundary was approximated to follow the interpolated pre-furrow shape (Figure 6b-e). To test for spatial correlations between wall tensile stress and microtubule localization, another FE model was constructed that followed the observed contour of the nascent furrow (Figure 6g). In this model, there was a very focused tensile stress maximum of ~440 MPa centered on the furrow apex at the junction of the outer periclinal and anticlinal walls. The high stress is due in part to the local high curvature at the apex, and this location also corresponded to the largest anticlinal microtubule peak along the entire segment (Figure 6h). Therefore, the biomechanics and cellular mechanisms of cell-autonomous furrow formation are indistinguishable from those of interdigitating pavement cells. Both processes can be explained by an “initiating” cell that uses a microtubule-cellulose biosynthesis module to convert cell wall stress patterns into a polarized growth response.

**Figure 6.**
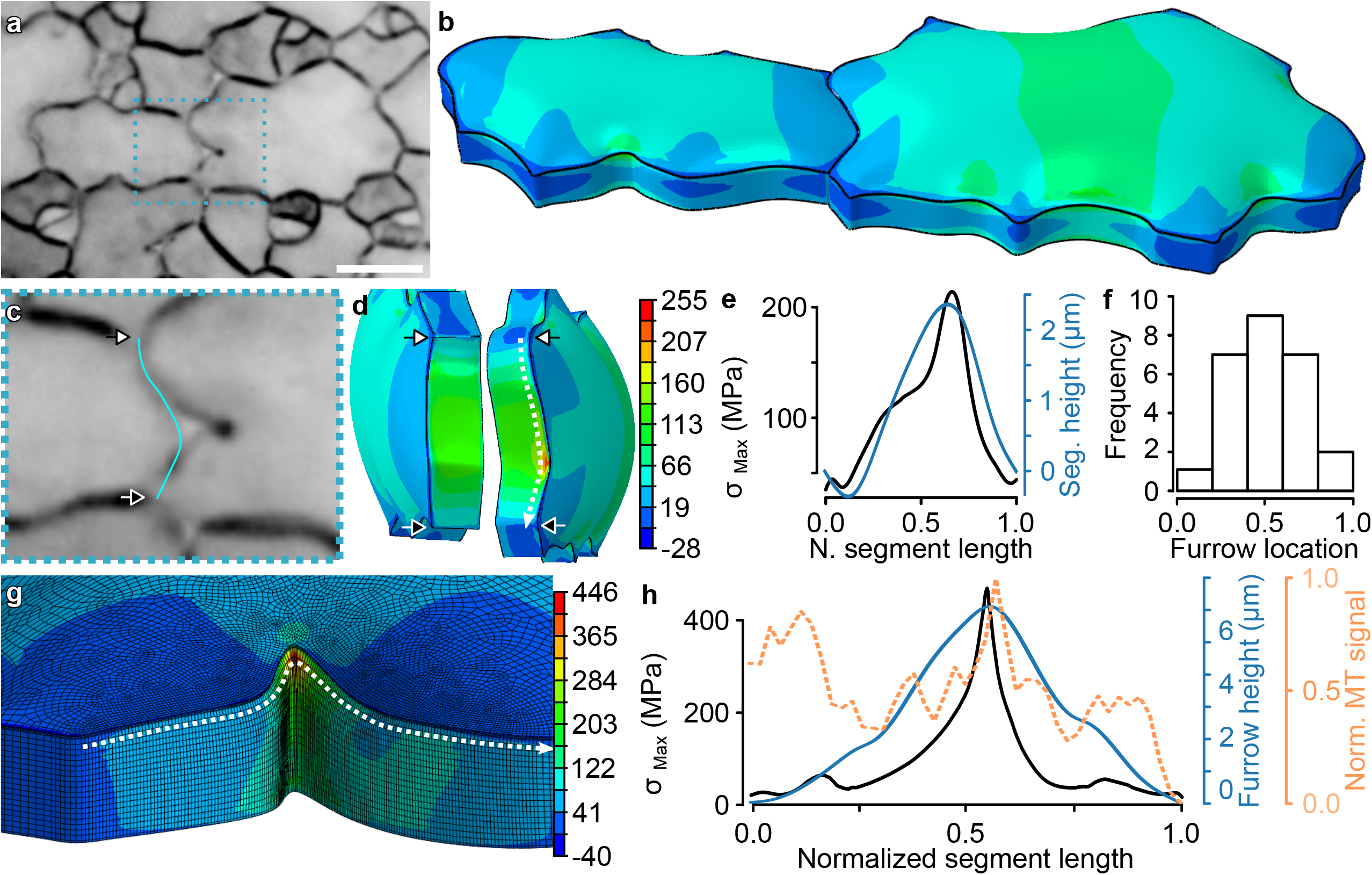
Cell wall stress dictates the location and direction of furrow formation. **a**, Face-on view of pectinase-treated cells displaying furrows. **b**, A finite element model of the estimated shape of the cell pair prior to furrow formation using the flanking cell contours shown in (**c**). **d**, Visualization of the anticlinal wall stress patterns of the furrowing cell (right) and its neighbor (left). **e**, Line plot of anticlinal wall stress and segment height relative to a baseline connecting the 3-way junctions. **f**, Population-level analysis of furrow location and normalized distance along the segment. **g**, Heatmap of cell wall stress of the furrowed cell in (**a**). **h**, Aligned peaks of cell wall stress (**g** – black line), microtubule (MT) signal (orange), and segment shape (blue) in a furrow. Scale bar = 10 μm.

To evaluate the potency of a microtubule-patterned cellulose-rich patch on cell shape change, an FE model was constructed based on a lobing cell pair with a 2μm wide anisotropic patch that mirrored the directional stiffness of cellulose fibers and spanned the anticlinal and outer-periclinal walls (Figure 7a-c). The anisotropic patch generated a region of elevated stress due to the transition in material properties (Figure 7d,e). After a single iteration of wall loading and relaxation, the anisotropic patch locally distorted the cell-cell interface to form a distinct convex feature compared to the isotropic control cell (Figure 7f,g). The presence of the anisotropic patch led to an additional deformation of more than 200 nm above that of the isotropic model, which is similar to the magnitude of stable features that were detected in time-lapse light microscopy experiments^15^. The anisotropic patch leads to a higher degree of localized and directional cell boundary deformation which supports the conclusions from the intact tissue and the cell-autonomous lobing system. This simulation demonstrates the ability of a microtubule-patterned cell wall heterogeneity at the micron scale to program a tissue-scale symmetry-breaking event.

**Figure 7.**
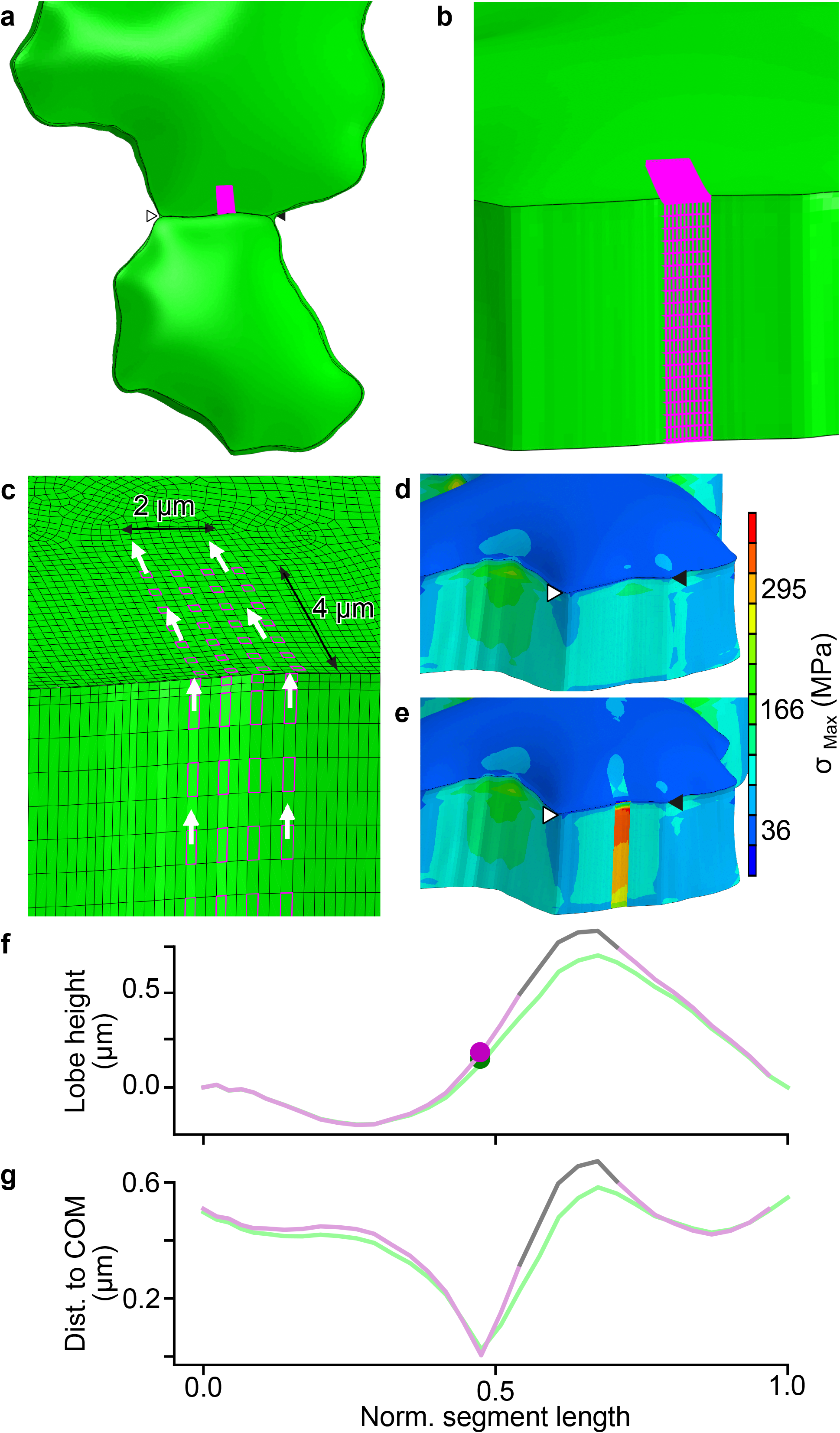
Localized microfibril alignment can drive lobe initiation: an FE analysis. **a**, Overview of a lobing pair of pavement cells. A transfacial patch of anisotropic material is located in one cell while the rest of the material in both cells is isotropic. **b**, The material in the connected anticlinal wall in the same cell is also defined as anisotropic. Note that no transition zone near the edges of the patch is used. **c**, The anisotropic patch has a width of 2μm, and the simulated MF direction is aligned with the arrows. **d**, Maximum principal stress map after pressurization and relaxation for the completely isotropic case; this result serves as a reference for the deformation. **e**, Stress map for the case which includes the anisotropic patch, which shows a stress concentration within the patch area. **f-g**, Comparison of the cell-cell interface geometry for the isotropic (reference - green) and anisotropic (magenta) models.

## Discussion

### Multiscale growth control

A major challenge in biology is to understand how the cytoplasm patterns the polarized growth of groups of cells to generate a tissue with specialized functions. Interdigitated growth in the leaf epidermis controls the expansion of thin, durable leaves^4,8^. Here, we used the Arabidopsis cotyledon development system to discover how cell wall tensile force and a specialized population of microtubules initiate interdigitated growth (Figure 8). Lobe formation is first detected as a local nanoscale ~300 nm warping^15^ that is driven by microtubule and cellulose microfibril-programmed anisotropy in a small patch of the cell wall in an initiating cell. These tiny features have a multiscale influence on morphogenesis because they become stabilized, and slow irreversible growth over a period of days morphs the feature into a well-defined lobe with features at ~10 - 50 μm scales^17,25,30^. This work defines the mechanism, timeline, and type of cell wall patterning that serves as a general model for polarized diffuse growth and provides a foundation to analyze epidermal morphogenesis across wide spatial scales.

**Figure 8.**
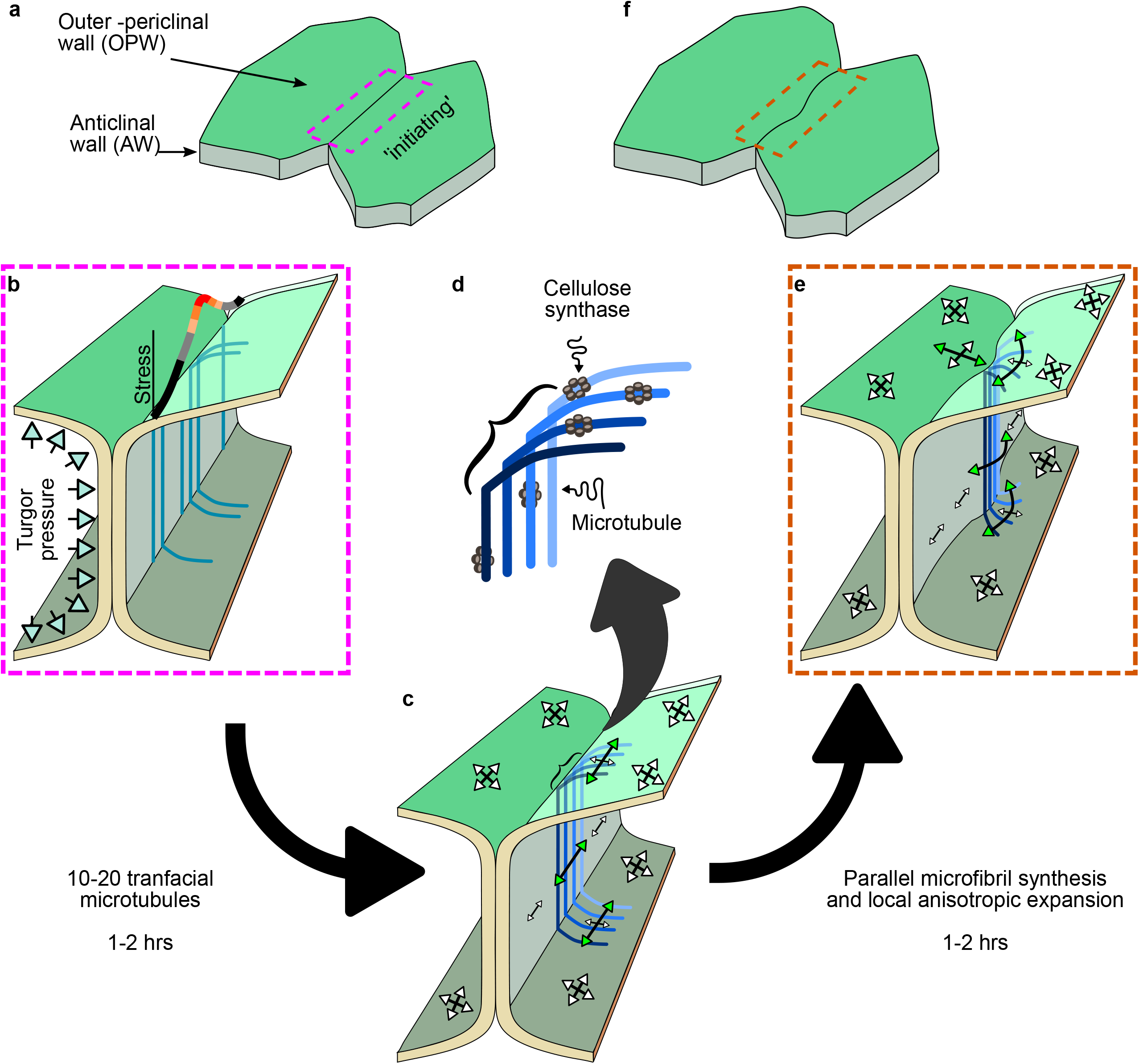
A general model for lobe morphogenesis and interdigitated growth. **a**, An example cell segment prior to lobe formation. **b**, Inset of (**a**), tensile stress is slightly elevated in the initiating cell due to geometric and wall composition variables. **c**, Microtubules are locally stabilized in response to the stress gradient. **d**, CESA complex motility in the plasma membrane is defined by local microtubule orientation, generating a small patch of the cell wall with anisotropic character. **e**, The initiating cells grow anisotropically in the region of the fiber-reinforced patch, and cell-cell adhesion and anisotropic expansion of the following cell in the direction of the anticlinal wall causes symmetry breaking. **f**, A recognizable new lobe is formed.

### The morphological potency of localized transfacial microtubules

Central to this conserved mode of tissue morphogenesis is a morphologically potent subset of transfacial microtubules (Figure 3; Supplementary Figure 4). No prior live cell study on lobe formation analyzed plasma-membrane and microtubule markers at a temporal resolution that would allow lobe initiation or the relevant microtubules to be detected^19,23^. Our multivariate live cell imaging pipeline captured microtubule organization and cell boundary changes along both the anticlinal and periclinal walls at 10 min intervals for 3.5 to 8 hrs. This enabled us to detect and quantify the strong enrichment of transfacial microtubules at cortical sites that would later generate lobes.

The central importance of the microtubule-cellulose system was supported by a series of mutant analyses and inhibitor studies. Only the mutation of *MOR1*, an essential microtubule-associated protein^38^, and *any1/cesa1* had clearly reduced lobe number phenotypes (Table 1). Other mutants affected lobe outgrowth or cell size, which are often misclassified as lobe initiation mutants based on manual lobe calls or dimensionless shape descriptors like circularity. Inhibitor studies also showed that compounds that disrupt microtubules and cellulose synthesis reduced lobe initiation, while pectinase treatment did not (Supplementary Table 1). Historically, microtubules have figured prominently in growth restriction models of lobe initiation^11,21^, and more recently additional pectin-based models for lobe initiation control have emerged^23,24^. In our assays, pectin mutants and pectinase treatment had the strongest effect on cell size, not lobe initiation. This phenotype likely reflects critical functions for pectin as the major matrix component of the wall that affects microfibril-microfibril interactions and cell wall stiffness. A complete loss of pectin would certainly be lethal. Short term pectinase treatment did not significantly reduce cell size (Supplementary Table 1), and the clearest effect of *qua2* and pectinase treatment in the context of tissue morphogenesis was to promote delamination and enable highly polarized cell-autonomous lobing (Figure 5; Figure 6; Table 1; Supplementary Table 1). This pectinase-induced invaginating growth required a functional microtubule-cellulose system (Figure 5f) and was mechanistically identical to the lobing of adherent cells (Figure 5 and Figure 6). The simplest explanation is that pectin has secondary functions during lobe initiation, by promoting cell-cell adhesion, and perhaps affecting the local modulus of initiating and/or following cells to enable lobe expansion. We emphasize that *qua2* and pectinase treatment does not completely remove pectin from the cell wall, and we are not questioning the general importance of this major cell wall matrix component.

The net vertical alignment of the microtubule is a stable feature of the anticlinal cell face of pavement cells (Supplementary Figure 5c,f and Belteton, et al.^15^). The resulting high degree of cellulose-encoded cell wall face-specific anisotropy restricts vertical expansion and is more permissive for expansion in the plane of the leaf surface. This organization appears to enable the horizontal component of tensile stress in the anticlinal wall to locally dictate growth rates (Figure 2f, Supplementary Figure 2l-o). Given that σ_H_ is on the order of 10 MPa acting on an anticlinal wall with a thickness of ~30 nm and a height of ~10 μm, it is estimated that a force of ~3 μN is needed to drive lateral expansion and increase the distance between oriented microtubules. Highly aligned microfibrils are also present in the anticlinal walls of epidermal cells of maize coleoptiles^47^, and it is possible that the microtubule-microfibril systems of this cell face are widely used to pattern polarized diffuse growth.

Transfacial microtubules have morphogenetic power in the context of lobing because they can efficiently align trajectories of CESA complexes and cellulose microfibrils across adjacent cell faces in a localized cortical domain (Figure 8). This model predicts that genetic and pharmacological effects of altered cellulose synthesis are indistinguishable from those of microtubules (Table 1; Supplementary Table 1; Supplementary Figure 3) and CESA trajectories align with microtubules in lobing pavement cells (Supplementary Figure 3r-t). Transfacial microtubules program the synthesis of a local patch of anisotropic cell wall. The end result is not growth restriction. Fiducial mark tracking along the anticlinal wall (Figure 1) and time-lapse analyses showed that lobe apices are actively growing features (Figure 1g, k; Supplementary Movie 3). An anisotropic patch of wall that mirrors fibers aligned along the anticlinal and periclinal surfaces of one cell has been hypothesized to promote anisotropic expansion parallel to the cell boundary and initiate lobe formation^13^. FE simulation reinforces the plausibility of this symmetry breaking mechanism and indicates that the proposed wall patterning in an initiating cell and the resulting growth behaviors can potently warp the cell boundary and initiate lobe formation (Figure 7).

Beyond a preferential enrichment of transfacial microtubules in the initiating cell, there was no unique microtubule behavior associated with lobe initiation. Both the degree and time interval of transfacial microtubule enrichment prior to lobe detection varied greatly among events (Figure 4e; Supplementary Figure 4t-y; Supplementary Figure 5o). This likely reflects the numerous parameters that influence the localization and morphogenetic power of microtubules. What is the nature of the cell wall stress profile along a particular cell-cell interface? How many CESA complexes were recruited during its lifetime? How many microfibrils were synthesized, of what length with what degree of coupling to the matrix? To what extent does microfibril-dependent CESA alignment locally operate as a positive feedback loop for anisotropic wall patterning^48^. What is the size, position, and degree of anisotropy in the cell wall patch? The stiffness of the outer periclinal wall of the adjacent “following” cell could also strongly affect lobe initiation. Clearly the notion that increased cell size drives lobe formation is too simplistic^22^. The time-lapse analysis of microtubule-dependent symmetry breaking shows that neither the site of lobe initiation nor its direction are determined solely by cell size or preexisting convex cell boundary (Supplementary Figure 5x-z). Further, the lobing outcome is not dictated by pre-existing cell wall properties, because in some instances bidirectional furrows formed at the same location in both cells (Figure 5c). Lobe formation is analogous to “a tug of war” between two mechanically coupled cells, in which small differences in tensile force tensors can bias transfacial microtubule occupancy, ultimately dictating the location and direction of lobe formation. Any parameter that affects tensile force locally can influence the polarity of the process. In two time-lapse experiments, there was a well-defined interval of ~2 hrs during which transfacial microtubules were enriched prior to lobe detection (Supplementary Figure 4u,y). These results indicate that relatively few (~10) localized transfacial microtubules can program a detectable morphogenetic output with a time lag of ~ 1 hr.

We would like to note that transfacial microtubules and sensitized growth control at the cell-cell interface may be a general feature of plant cell and tissue morphogenesis. Diffuse growth is assumed to be uniform across the cell surface. However, when densely distributed particles are used as fiducial marks, even in relatively simple shaped hypocotyl epidermal cells, considerable subcellular growth heterogeneity was detected^25^. Hypocotyl cells adopt several distinct microtubule configurations on different cell faces^49^, and in general the subset of microtubules and the subcellular patterns of growth that dictate shape change are not known.

### Cell wall stress is a multiscale upstream patterning element

Lobing follows a minimum spacing rule^40,43^, and the underlying cellular patterning mechanism for lobe formation has been controversial^14,15,21,50^. Here we used FE models of pavement cell clusters to simulate the magnitude and direction of cell wall stress patterns quantitatively and test for correlations with microtubule behaviors. We show that microtubules were correlated with stress in 3 different ways. First, microtubules are depleted at 3-way junctions^15^, and wall tensile forces are often reduced near the mechanically stabilized 3-way cell wall (Figure 2c, Supplementary Figure 2; Figure 4g-j; Figure 6d,e). Second, microtubules along the anticlinal walls are highly aligned perpendicular to the leaf surface (Figure 3b,i; Supplementary Figure 5c,f; and see Belteton, et al.^15^) as are the maximum principal cell wall stresses (Figure 2e; Supplementary Figure 5b,e). They also have potential to globally align the microtubule-microfibril systems along the anticlinal face and influence local microtubule alignment of the outer cell face when subsets polymerize across the cell-face boundary. This proposed control system could promote a planar bias to cell expansion in any epidermal cell type with this organization^47^, and in pavement cells the orientations of these stress tensors likely contribute to differential growth rates along the anticlinal segments (Figure 2f, Supplementary Figure 2l-o). As σ_H_ and growth rates are elevated in subsets of lobes (Supplementary Figure 2m), lobe features and resulting increases in σ_H_ could enable increased growth rates from cell to organ scales. Third, there were local anticlinal and periclinal wall tensile stress peaks that correlated with the location of transfacial microtubule enrichments and subsequent lobe initiation sites (Figure 4i,j; Supplementary Figure 5s,w). These stress profiles are estimates of those that exist in real cells because the wall was treated as a homogeneous material, but nonetheless they consistently point to the ability of the microtubule system to sense subcellular gradients of tensile stress.

Wall stress patterns were strongly altered in pectinase-treated cells with broad delaminations and cell invaginations. In this cell autonomous lobing system wall stress maxima predicted the sites of delamination. Delamination was not an elastic response because it required polarized growth and an intact microtubule-cellulose system (Figure 5f). Asymmetric delamination had an apical curvature with a highly focused cell wall stress maximum. The strong and consistent concentration of microtubules at the furrow apex, and the active invagination of furrows that developed in a microtubule- and cellulose-dependent manner also indicate that local wall stress gradients are sensed to pattern polarized diffuse growth. In the context of an intact tissue, outer periclinal wall tensile forces near the cell boundary tend to be reduced in concave regions and elevated in convex regions and around the mid-regions of extended straight domains (Supplementary Figure 2k,o; convex – cyan arrowheads, concave – orange arrowheads). The coupling of these tensile stress patterns to the microtubule-microfibril system could explain the spacing rules of lobe formation.

Our TEM analysis showed that the morphology and microtubule localization in pectinase-induced furrows (Figure 5e) were identical to those of highly furrowed leaf mesophyll cells with a terminal bulb^45^. We believe that we have described a general mechanism of lobe formation. In specialized mesophyll cells, this subcellular trait is linked to more efficient CO_2_ transport to chloroplasts and has clear importance in crop productivity^2,9^. A key challenge now is to determine how cortical microtubules sense the large and oriented tensile forces in the cell wall without being ripped apart.

Cell wall stresses serve as useful multiscale patterning elements during tissue morphogenesis. Tensile forces are concentrated at the interface where the unpaired outer periclinal walls pull upward on anticlinal walls with stresses that reflect cell autonomous parameters like cell size and local height. Tissue-level parameters like the length and local curvature of the shared anticlinal wall also shape subcellular stress gradients that are decoded by the microtubule system. Along the anticlinal wall, the distance from and angles among cell walls at 3-way junctions^51^ may also affect stress distributions. Tensile stress is sensitive to the material property gradients in the cell wall and varies inversely with cell wall thickness^30^. Wall stress patterns may also reflect the local growth status, because within a pavement cell, growth rates within a segment often vary by factors of 2 to 7 (Figure 1f and Supplementary Figure 2j), and wall thickness varies by a factor of ~2.5 ^15^ due to imperfect maintenance of wall thickness during cell expansion. Interestingly some of the largest microtubule bundles may be localized to the thinnest regions of the anticlinal wall (see Figure S13 C and D Belteton, et al.^15^), and this may reflect a general wall stress sensing mechanism that prevents cell rupture. Our analyses indicate that cell wall stress patterns influence microtubule array organization at cellular scales, and the microtubule-cellulose system uses cell wall stress to reliably convert numerous geometry and physical parameters into a patterned morphogenetic response.

### Conclusions and Future Perspectives

Our integrated experimental data and FE modeling analyses point to the central importance of the shared anticlinal wall^16^ and the periclinal-anticlinal wall interface during mechanical signaling and polarized morphogenesis. The shared wall integrates stress patterns, growth status, and determines the local rates and direction of growth. These new insights into pavement cell shape control provide a new way to integrate realistic cellular growth mechanisms into models that seek to explain how planar organs develop^52-55^. However, at the moment, we do not know how stress- and microtubule-mediated growth patterns might scale to affect tissue-sector or organ morphology. The imaging locations here were primarily within the basal mid-blade of 1.5 DAG cotyledons, and the contributions of these cells and lobes to organ-scale shape change is not known. Organ-level leaf patterning is influenced by the patterned expression of combinations of proteins that somehow promote or limit growth^53,55^. Further analyses grounded in plausible FE growth models can reveal how morphogens, cell wall stress, and cellular growth machineries interact to sculpt the macroscopic traits of leaves.

## Acknowledgments

We thank David Jackson (Cold Spring Harbor Laboratory, Ithaca, NY) for his generous gift of PDLP3-GFP transgenic line and Ying Gu (Pennsylvania State University, University Park, PA) for her generous gift of the *csi1-3* mutant and *csi1-3*; YFP-CesA6; RFP-TUA5 transgenic line. This material is based upon work supported by the National Science Foundation under Grant No. 1715544 to DBS.

## Author contributions

Conceptualization, D.B.S., S.B.; Methodology, S.B. and W.L.; Investigation, S.B., W.L., M.Y., M.Q., M.S., M.M., and F.A.; Writing - Original Draft, S.B. and D.B.S.; Writing - Reviewing & Editing, S.B., W.L., M.Y., M.Q., M.S., M.M., F.A., J.T., and D.B.S.; Project Administration, D.B.S.; Funding Acquisition, D.B.S. and J.A.T.

