## supplemental data for "Real-time conversion of tissue-scale mechanical forces into an interdigitated growth pattern"

**Supplemental Figure Legends**

Supplementary Figure 1. Growth dependent separation of PDLP puncta pairs along an example segment. **a**, Using the plasma-membrane marker segments were traced at the anticlinal/outer-periclinal junction. **b-c**, Segment straightening followed by re-slicing produces a face-view of the anticlinal wall. **d**, Non-overlapping PDLPs including manually labeled 3-way-junctions were tracked over time. **e**, PDLP-pair distance is used to obtain sub-segmental growth rates from linear fit models over the time course. **f**, Sub-segmental elemental growth rates mapped onto the initial segment shape. PDLPs are stable features in the anticlinal wall and their displacement reflects localized diffuse growth. **g-i**, Cold-treated PDLP3:GFP; PIP2:mCherry seedlings were imaged at 4-hour intervals. **j-k**, Normalized signal intensity plots and overlapping peaks from PDLP3 particles of the two segments (**g-i**) at all 3 time-points. **l**, Boxplots of measured sub-segmental growth rates of cold-treated (CT) and room temperature (RT) grown seedlings. Sub-segmental growth rates from lobing segments. **m-o**, Plots of normalized segment length at lobe detection (gray outline) and final (black outline) time-points color coded for normalized percent growth rate per hour. Cyan and green arrowheads mark the location of newly and established lobes respectively (Top panel). Perimeter to center of mass plots highlight warping regions between lobe detection and final timepoint.

Supplementary Figure 2. **a**, Microscopic image of the cells which were used to study the effect of the partial geometry of cells for the stress response under turgor pressure. The star shows the location of the maximum deflection of the periclinal wall for Cell 2. **b**, 3D finite element (FE) model of the two complete cells and the distribution of maximum principal stress (Max PS) on both anticlinal walls (AW). **c**, Distribution of the mean of the Max PS from the upper $1/8^{th}$ of the AW for both cells for three example results of partial cells in terms of the area used in the analysis. **d**, Maximum of the absolute value of the percentage difference between mean Max PS distributions in the upper $1/8^{th}$ of the AW in each simulation relative to using the complete geometry of cells. It indicates that the FE analysis can use a partial cell. If the area includes the maximum deflection the distribution of the Max PS does not change substantially and the magnitude is reduced by $< 10\%$. **e**, Comparison of the average of Max PS from the different upper areas of materials in the AW. It turns out that the mean of Max PS of the different upper areas in the walls has the same distribution pattern but varies in magnitude. **f**, Optical image of the cells were used to study the correlation of the Max PS and the growth rate of the cell-cell interfaces. **g**, Cell 2 from the measurement is defined partially, but the maximum deflection area can be seen in the image. **h-i**, Comparison of maximum deflection of periclinal wall for Cell 2 in (**f-g**) from the simulations using different remaining areas of the cell. The results of (**i**) is very consistent with the optical measurement in (**g**). **j-l**, Like the study described in (**f**), four other segments of cells were used for the comparison, and the results show that the horizontal component of Max PS along the segment length has a strong correlation with the growth rate of the associated wall. The scale bar for all = 10 μm.

Supplementary Figure 3. The microtubule and cellulose systems are required for lobe formation. Mature pavement cell shapes, magenta outline, were analyzed using LobeFinder. Mutants affecting the microtubule- (**b**, **c**, **i-k**), cellulose- (**d**, **e**), and pectinase- (**f**), and secretion (**g**) were compared to wild-type (**a**, i). The temperature-sensitive mutant *mor1-1* was tested at permissive (**h**) and restrictive temperatures (**j**) and compared to its wild-type counterpart (**i**). Effects of 0.2%

pectinase (**m**), 1μM oryzalin (**n**), and 5nM Isoxaben (**o**) on cell shape from 1 to 2 DAG compared to a buffer treated control (**l**). A combination of 0.2% Pectinase with a reduced concentration of Oryzalin [1μM] (**p**) and the combination of 0.2% Pectinase and 5nM Isoxaben (**q**) both completely halted the production on new lobes (Table S1 – for results). Cellulose synthase complex marked with YFP:CesA6 strongly colocalizes with cortical microtubules in early stage pavement cells. **s**, 8-min projection of 15-sec intervals, single plane time course shows how CesA tracks resemble and co-localize (**t** – white arrows) with cortical microtubules, mCherry:TuA5 (**r**). Evidence of colocalization between cellulose synthase complex marked with CesA3:GFP and cortical microtubules at the anticlinal wall. **u**, Top-down view of segment resliced. Dotted line shows shape and direction of segmentation. White arrows on resliced images are CesA complexes that localized to anticlinal microtubules. The colocalization of cellulose and microtubules was not affected in 8-minute projection of 15-sec intervals, single plane time-lapses, in the *csi1* mutant background where CesA tracks (YFP:CesA6 - **w**) resemble and co-localized (**x** – white arrows) with cortical microtubules, RFP:TUA5 (**v**). Resliced views of the anticlinal wall also show evidence of cellulose and microtubule co-localization in the *csi1* mutant. CesA colocalization with anticlinal microtubules is evident in the *csi1* mutant background. **y**, Top-down view of segment resliced with dotted line showing the shape and direction of segmentation. White arrows on resliced images on the right are CesA complexes that localize to anticlinal microtubules. The anticlinal wall at the apex remains straight during lobe formation. **z**, Snapshots of a lobing segment before, (aa) during, (**bb**-**cc**) and after lobe formation. **dd**, Cross-sectional views of the anticlinal wall at the lobe apex from locations specified in (**z**-**cc**). **ee**, Histogram of anticlinal wall tilt showing how the vast majority of measure wall regions were completely perpendicular to the leaf surface. Scale bars = 50μm (**a**-**q**), 25μm (**k**-**q**), 5μm (**r**-**dd**).

Supplementary Figure 4. Analysis of microtubule turnover in time series data. Pairwise correlation results between the initial time-point and each subsequent time-point (blue line), a sliding window of paired consecutive frames over the time course (green dashed line), and each time-point to a randomly selected time-point (magenta dotted line). The random correlation results average (gray dotted line) is the background level for microtubule correlations. **a**-**b**, 15-sec interval time-lapses. **c**-**d**, 1-min interval time-lapses. **e**-**f**, 5-minute interval time-lapses. Each sampling regime had a similar pattern of correlation decay from the first time-point to background levels after ~10 min. Microtubule orientation in ROIs in opposing cells that undergo lobe initiation does not show detectable differences in alignment and coherence. **g**-**i**, Snapshots of plasma-membrane (PIP2:mCherry) of a lobing segment, yellow boxes are ROIs used to monitor microtubule behaviors in opposing cells lobe centered at the future lobe region. **j**-**l**, Snapshots of the microtubules (GFP:TUB6). Microtubules orientation and coherency results using red ovals whose tilt reflects the orientation and their aspect ratios reflect the coherency of the microtubules at the time point shown. **m**-**n**, Scatterplot of microtubules orientation (green) and microtubule coherency (purple) in cell 1 (**g** -top) and cell 2 (**g** - bottom) as a function of time before (< time-point 19) and after (≥ time-point 19) lobe detection. Distribution of manually identified transfacial microtubules in lobing segments. **o**-**s**, Top - segment shape before (blue) and after (orange) lobe detection. Yellow arrowheads are existing lobes. Cyan arrowheads are newly formed lobes. The top histogram (green) depicts transfacial microtubules at the future convex cell at the lobe initiation site. The bottom histogram (magenta) depicts transfacial microtubules of the future concave cell at the lobe initiation site. Time-lapsed analysis of the occurrence and location of transfacial microtubules in six different lobing segments. **t**-**y**, The number of transfacial microtubules in the lobing cortical domain in the future convex (green) and future concave (magenta) cells. Scale bar = 5µm.

Supplementary Figure 5. Semi-automated analysis of accumulated microtubule signals detects local microtubule enrichment at the future lobe initiation site. (**a**, **d**) Heatmap of σ-max values on the anticlinal wall for two lobing segments. (**b**, **e**) Density plots of stress alignment at the anticlinal wall for cell 1 (green) and cell 2 (magenta). (**c**, **f**) Face-view of cortical microtubules at the anticlinal wall for segment depicted on A and D respectively. **g**-**i**, Time-lapse images of the plasma-membrane, (PIP2:mCherry - magenta) and microtubules (GFP:TUB6 - green) of a lobing segment that was captured for > 4 hours before the lobe was detected. **j**-**l**, Outline of the segment shape for segment corresponding to the live-cell images above where the lobe initiation cortical domain is marked using cyan vertical lines and lobes peak locations are marked by a black vertical line. **m**, Density plot of manually scored transfacial microtubules where cell 1 has a peak at the future convex side of a lobe before its detection. **n**, The accumulated microtubule signal method can detect enriched microtubule signal ($Enrichement = 1 - \frac{Area\ under\ curve_{Cell1}}{Area\ under\ curve_{Cell2}} * 100$) in cell1 within the future convex domain. **o**, Summary of outer periclinal microtubule enrichment in future convex cells measured from 11 independent lobing events. Boxplots show the distribution of microtubule enrichment of future lobe convex domain prior to lobe detection. The number of time-points at 10 min intervals is listed above each boxplot. Example images of two additional lobing pairs that could be converted to FE models and tested for MT and tensile stress correlations at sites of microtubule enrichment and lobe formation. (**p**, **t**) Summed microtubule signal projected images of segments before lobe detection. (**q**, **u**) Enrichment of microtubules at the future convex side of a lobe as a function of time before lobe detection. (**r**, **v**) Line plots of FE $\sigma_{Max}$ predictions along the anticlinal wall of cell 1 (green) and cell 2 (magenta), microtubule accumulated signals along the shared anticlinal wall (black dashed line). (**s**, **w**) Line plots of FE $\sigma_{Max}$ predictions along the outer periclinal wall of cell 1 (green) and cell 2 (magenta), microtubule accumulated signals outer periclinal wall of cell 1 (green dashed line) and cell 2 (magenta dashed line). Symmetry breaking events are not defined by a pre-existing segment shape. **x**-**z**, Snapshots of eight-hour time-lapse with plasma-membrane (PIP2:mCherry - magenta) and microtubule (GFP:TUB6 - green) markers of a segment that has a symmetry-breaking event against the established bulging shape. (**aa** - top) Accumulated microtubule signal plot and level of enrichment within the future convex cortical region (cell 1) (**aa** - bottom) Accumulated microtubule signal plot (cell 2). Scale bars = 5µm for **c**, **d**, **g**-**i**; 10µm for **x**-**z**.

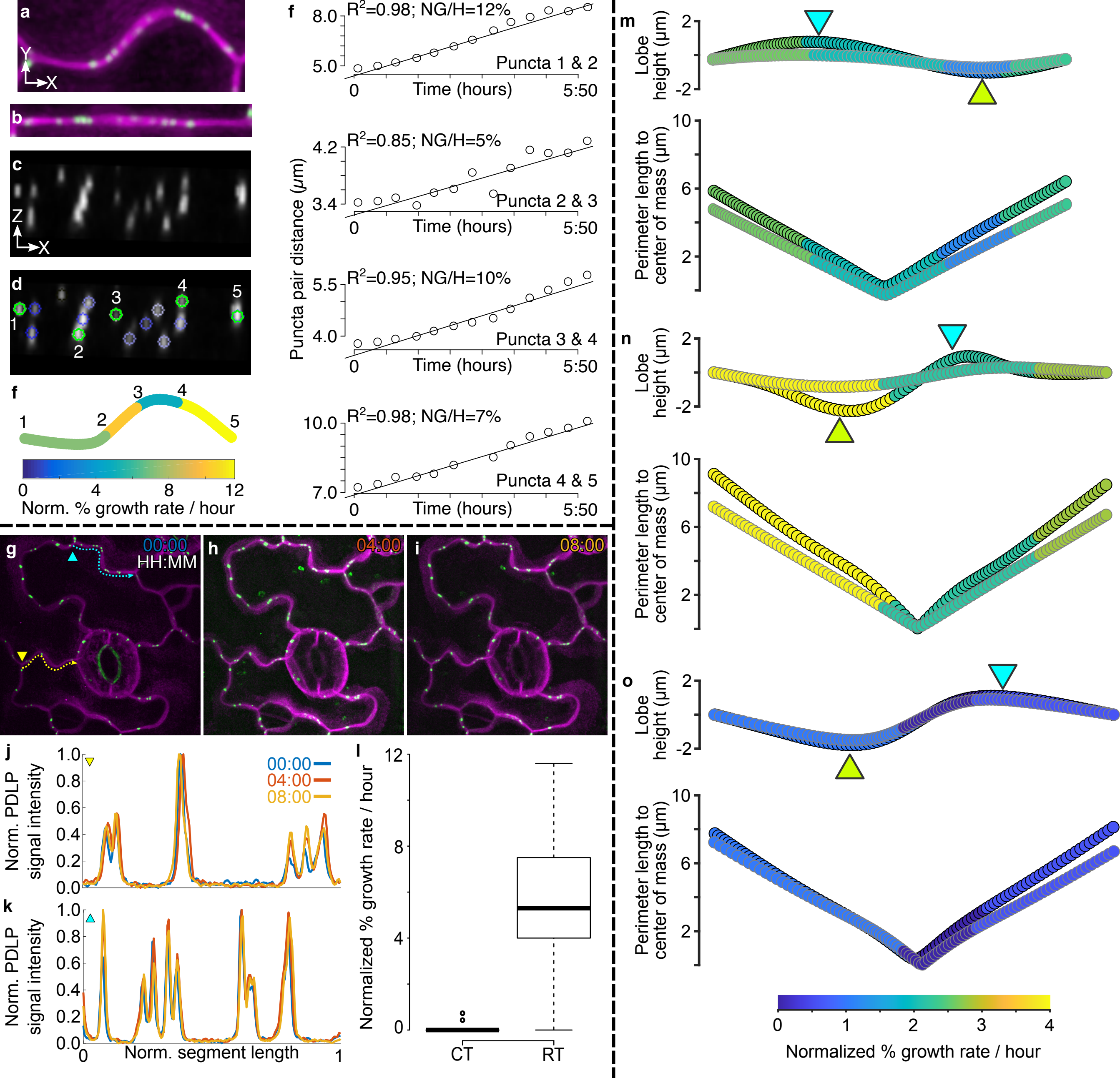

a
Y
X
b
c
Z
X
d
1
2
3
4
5
f
1
2
3
4
5
0
4
8
12
Norm. % growth rate / hour
f
Puncta pair distance (μm)
R²=0.98; NG/H=12%
Puncta 1 & 2
R²=0.85; NG/H=5%
Puncta 2 & 3
R²=0.95; NG/H=10%
Puncta 3 & 4
R²=0.98; NG/H=7%
Puncta 4 & 5
Time (hours)
0
5:50
g
00:00
HH:MM
h
04:00
i
08:00
j
Norm. PDLP signal intensity
00:00
04:00
08:00
k
Norm. segment length
l
Normalized % growth rate / hour
CT
RT
m
Lobe height (μm)
Perimeter length to center of mass (μm)
n
o
Normalized % growth rate / hour

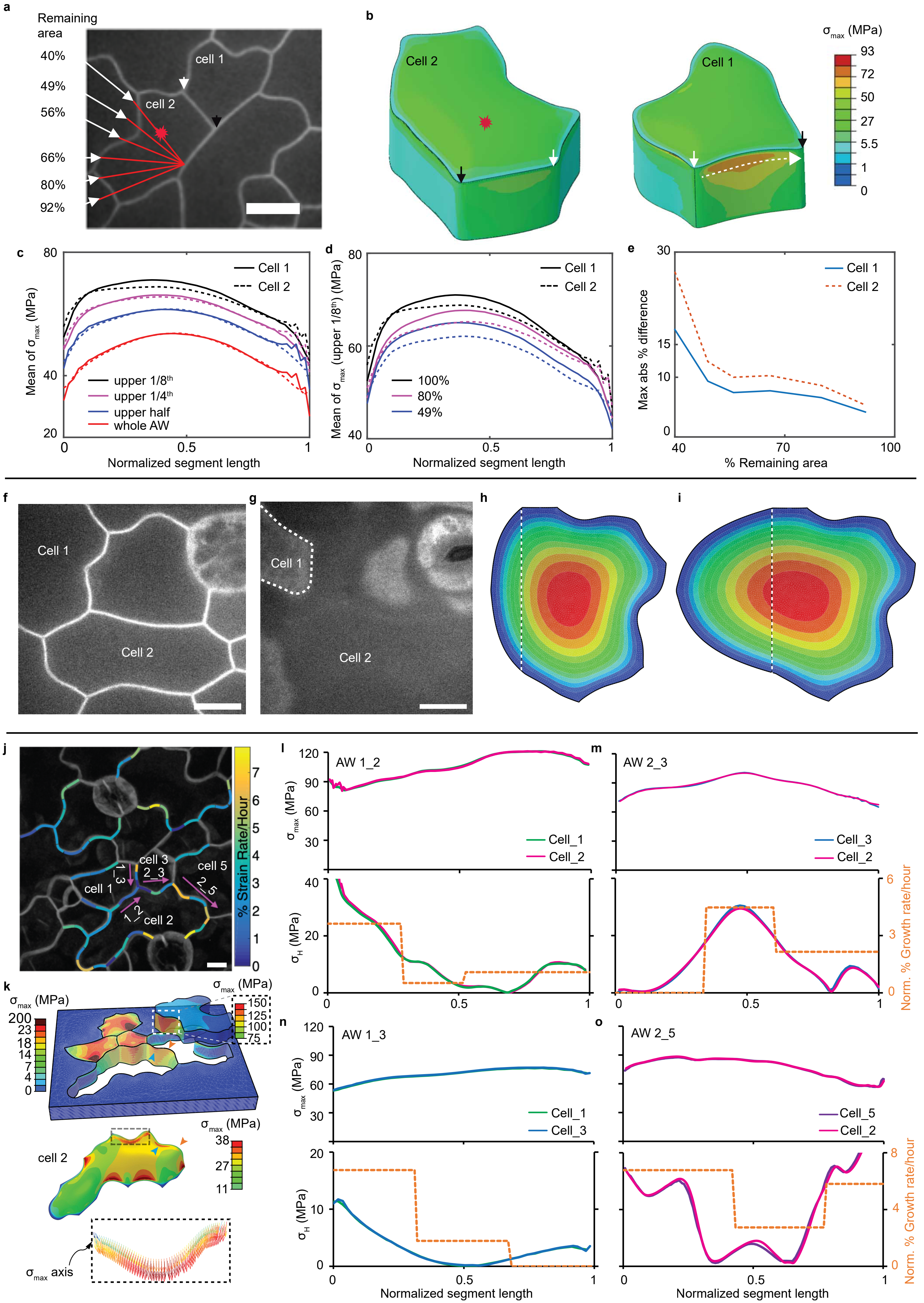
a
Remaining area
40%
49%
56%
66%
80%
92%
cell 1
cell 2
b
Cell 2
Cell 1
σmax (MPa)
93
72
50
27
5.5
1
0
c
Mean of σmax (MPa)
Normalized segment length
Cell 1
Cell 2
upper 1/8th
upper 1/4th
upper half
whole AW
d
Mean of σmax (upper 1/8th) (MPa)
Normalized segment length
100%
80%
49%
e
Max abs % difference
% Remaining area
f
Cell 1
Cell 2
g
Cell 1
Cell 2
h
i
j
% Strain Rate/Hour
cell 1
cell 2
cell 3
cell 5
1_3
2_3
1_2
2_5
k
σmax (MPa)
cell 2
σmax axis
l
AW 1_2
σmax (MPa)
σH (MPa)
Cell_1
Cell_2
m
AW 2_3
Cell_3
Cell_2
Norm. % Growth rate/hour
n
AW 1_3
Cell_1
Cell_3
o
AW 2_5
Cell_5
Cell_2
Normalized segment length

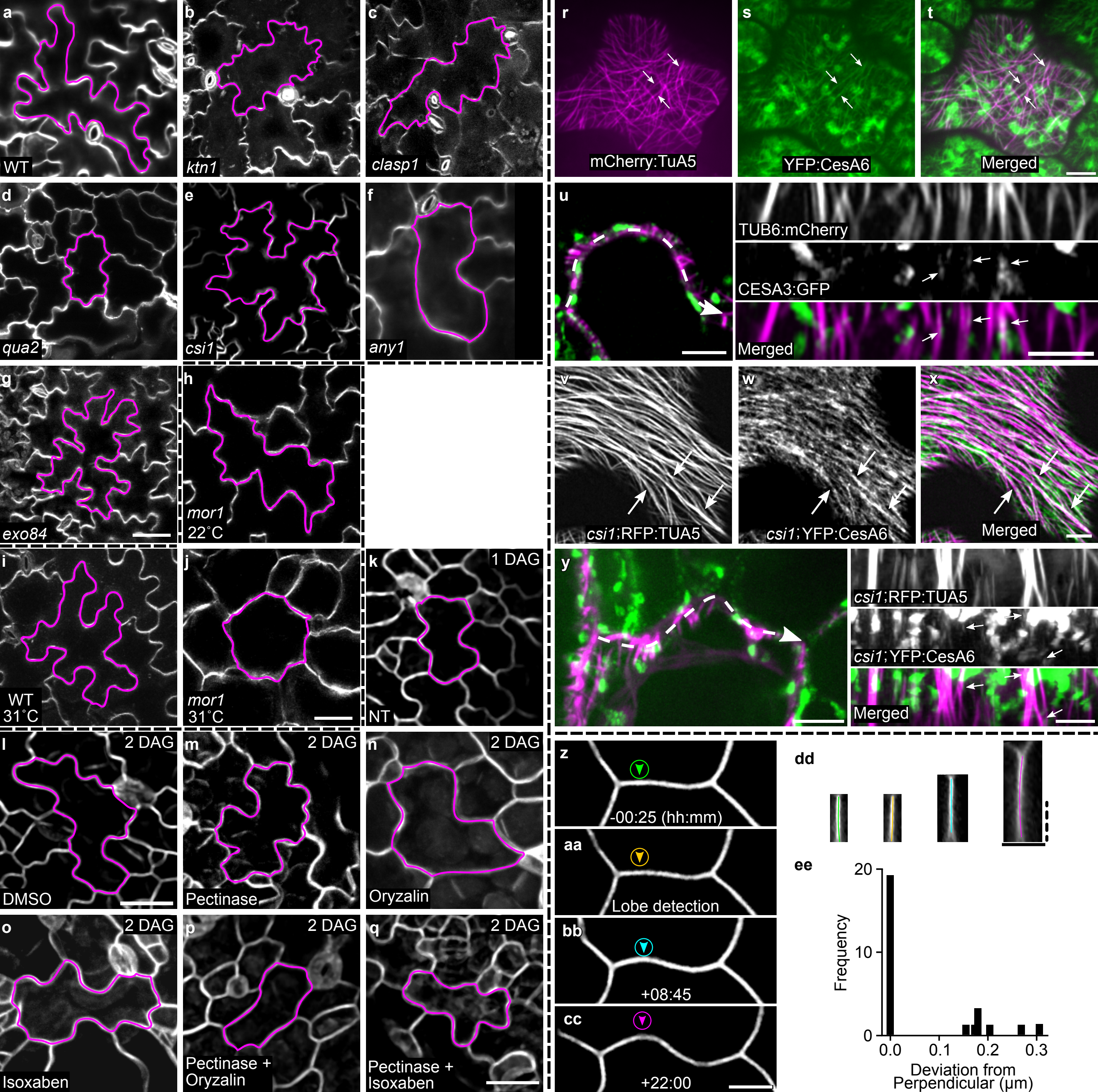

a WT
b ktn1
c clasp1
d qua2
e csi1
f any1
g exo84
h mor1 22˚C
i WT 31˚C
j mor1 31˚C
k 1 DAG NT
l 2 DAG DMSO
m 2 DAG Pectinase
n 2 DAG Oryzalin
o 2 DAG Isoxaben
p 2 DAG Pectinase + Oryzalin
q 2 DAG Pectinase + Isoxaben
r mCherry:TuA5
s YFP:CesA6
t Merged
u TUB6:mCherry
CESA3:GFP
Merged
v csi1;RFP:TUA5
w csi1;YFP:CesA6
x Merged
y csi1;RFP:TUA5
csi1;YFP:CesA6
Merged
z -00:25 (hh:mm)
aa Lobe detection
bb +08:45
cc +22:00
dd
ee
Frequency
Deviation from Perpendicular (µm)
0.0
0.1
0.2
0.3
0
10
20

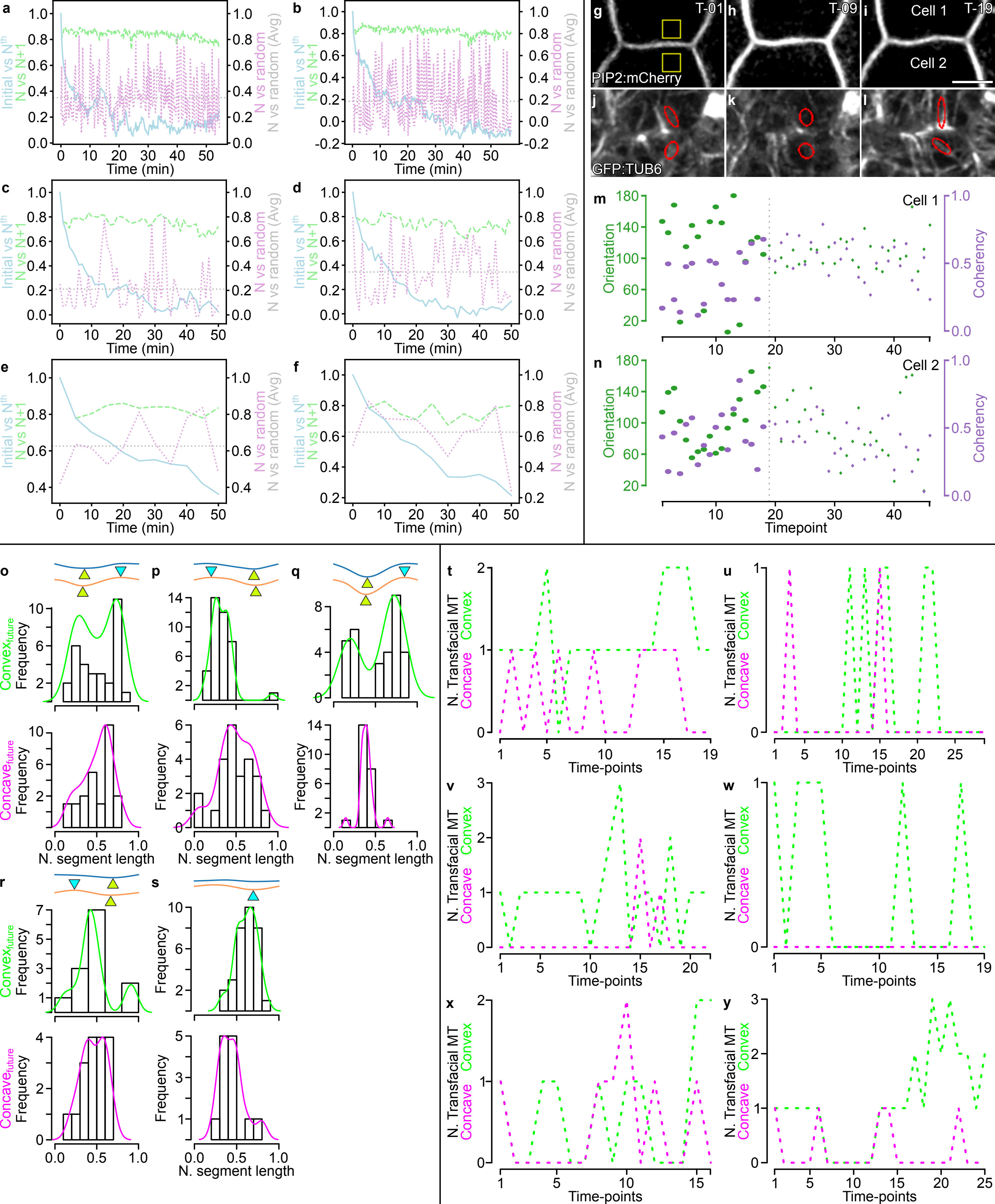
a
b
c
d
e
f
Initial vs N^th
N vs N+1
N vs random
N vs random (Avg)
Time (min)
g
h
i
T-01
T-09
T-19
Cell 1
Cell 2
PIP2:mCherry
j
k
l
GFP:TUB6
m
n
Orientation
Coherency
Timepoint
o
p
q
r
s
Convex_future
Concave_future
Frequency
N. segment length
t
u
v
w
x
y
N. Transfacial MT
Concave
Convex
Time-points

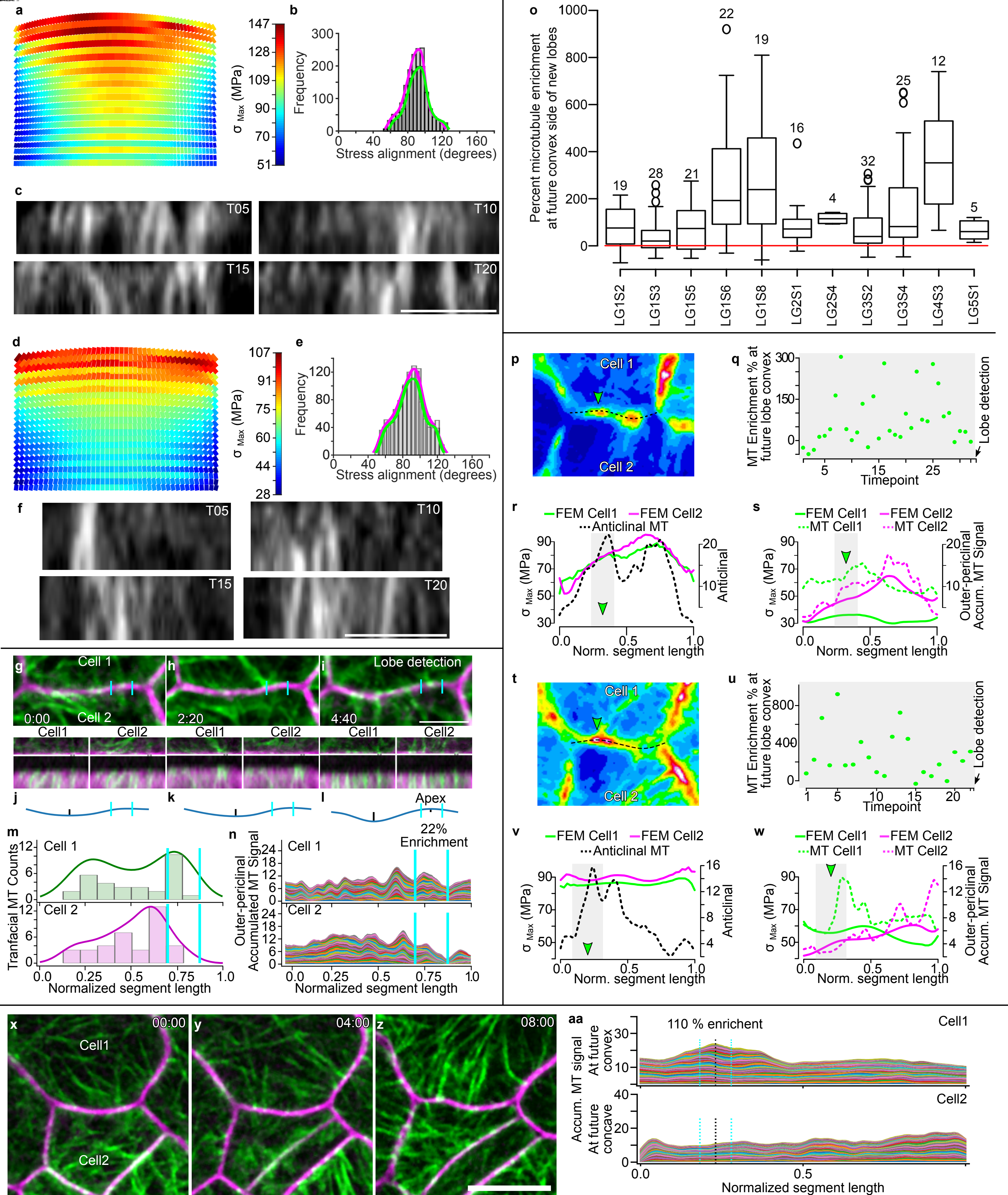

a
b
c
d
e
f
g
h
i
j
k
l
m
n
o
p
q
r
s
t
u
v
w
x
y
z
aa
σ Max (MPa)
Frequency
Stress alignment (degrees)
T05
T10
T15
T20
Percent microtubule enrichment at future convex side of new lobes
LG1S2
LG1S3
LG1S5
LG1S6
LG1S8
LG2S1
LG2S4
LG3S2
LG3S4
LG4S3
LG5S1
Cell 1
Cell 2
Cell1
Cell2
0:00
2:20
4:40
Lobe detection
Apex
22% Enrichment
Tranfacial MT Counts
Normalized segment length
Outer-periclinal Accumulated MT Signal
MT Enrichment % at future lobe convex
Timepoint
FEM Cell1
FEM Cell2
Anticlinal MT
MT Cell1
MT Cell2
Norm. segment length
Anticlinal
Outer-periclinal Accum. MT Signal
00:00
04:00
08:00
110 % enrichent
Accum. MT signal
At future convex
At future concave

Supplementary Table 1. Microtubules and cellulose synthesis are required for normal lobe initiation.

| | | | Area (μm$^2$) | Circularity | Lobe Number | Lobes/Area*1000 |
|---|---|---|---|---|---|---|
| DAG | Treatment | N | mean ± SD | mean ± SD | mean ± SD | mean ± SD |
| 1 | NT | 40 | 634 ± 163 | 0.62 ± 0. 08 | 7 ± 2 | 11 ± 3 |
| | NT | 120 | 1254 ± 484* | 0.45 ± 0.09* | 11 ± 2 | 8 ± 2* |
| | DMSO | 60 | 1717 ± 531*◊ | 0.41 ± 0.08*◊ | 12 ± 2* | 7 ± 2*◊ |
| 2 | Oryzalin | 60 | 1852 ± 533*◊ | 0.62 ± 0.10◊ | 8 ± 2*◊ | 5 ± 1*◊ |
| | Isoxaben | 101 | 1450 ± 443* | 0.51 ± 0.08*◊ | 9 ± 2*◊ | 7 ± 2*◊ |
| | Pectinase | 60 | 1332 ± 334* | 0.47 ± 0.09* | 11 ± 2* | 8 ± 2* |

(*), significantly different than 1DAG NT, and (◊), significantly different than 2 DAG NT or for oryzalin different than 2 DAG DMSO control, Wilcox-Whitney test ($p < 0.05$)

**Plant Material and Growth Conditions**
Arabidopsis thaliana seedlings were grown at 22˚C on ½-strength Murashige and Skoog medium with 1% Sucrose (w/v) and 0.8% (w/v) Bacto-agar under continuous illumination. For the chemical treatments, [1µM] oryzalin (Dow AgroSciences) , [5nM] isoxaben (Sigma), 0.1% (v/v) DMSO (Sigma), and 0.2% (w/v) pectinase from Aspergillus niger (Sigma), the seeds were transferred and completely submerged in liquid media (which omits the Bacto-agar) after germination under continuous illumination. For low-light chemical treatment, the seedlings were grown on plain media until 1 DAG and then transferred to plates with the chemical compound dissolved onto the media which were placed vertically at low light (5µmol m−2 sec−1) at room temperature. Arabidopsis ecotype Columbia-0 was used as the wild type. Mutants lines were described previously: *any1*[1], *qua2-1*[2], *exo84b*[3], *clasp1*[4], *ktn1-2*[5], *dis2-1*[6], *zwi3*[7], *csi1-3*[8]. PDLP3::PDLP3:GFP[9] was crossed to PIP2:mCherry line[10]. PIP2:mCherry; TuB6:GFP double-tagged line[11]. mCherry:Tua5; YFP:CesA6[12].

**Imaging and Analysis of Cotyledon Pavement Cell Shape: Population-Level studies**
Whole seedlings were stained using 1µM FM4-64 (Invitrogen) for 30 min at 4˚C. Cotyledons were dissected and mounted into a Vaseline-lined chamber slide and imaged using a Bio-Rad 2100 laser-scanning confocal microscope mounted on a Nikon Eclipse E800 stand. Samples were excited with a 488-nm laser, and the fluorescence signal was collected using a 490-nm long-pass dichroic and a 500- to 550-nm band-pass emission filter using a 20x 0.5 NA objective. Image fields were obtained from the apical 1/3 of the cotyledon. Representative complete pavement cells not part of the stomata cell linage were traced manually using the polygon tool in Fiji and the traces were splined using the line-tool option. Pavement cell traces were analyzed using LobeFinder[13] and statistical analyses were done in RStudio.

**CesA colocalization with microtubules imaging**
1 DAG seedlings were mounted on a Vaseline-lined chamber slice and imaged using a 100x Plan-APO 1.46 NA oil-immersion objective and images were acquired using a spinning disk CSU-10 confocal head (Yokogawa Electric) mounted on a Zeiss Observer.Z1 inverted microscope controlled using SlideBook software (Intelligent Imaging Innovation). YFP and mCherry were excited by 515 and 561 lasers, respectively. A single optical section of the outer-periclinal wall was selected using the mCherry line. 15 second interval imaging was conducted for 8 minutes keeping the region of interest in focus by using the mCherry line. Qualitative time-projected images were constructed after using the bleach correction macro with the histogram matching method[14].

**Center of mass measurements and subsegmental growth rate analysis**
Confocal Microscopy and Time-lapse Imaging of Lobe Initiation; 1.5-DAG whole seedlings were mounted in our in-house long-term chamber system as previously described[15]. Confocal fluorescence microscopy was performed using a 100x Plan-APO 1.46 NA oil-immersion objective and images were acquired using a spinning disk CSU-10 confocal head (Yokogawa Electric) mounted on a Zeiss Oberver.Z1 inverted microscope controlled using Slidebook software (Intelligent Imaging Innovation). GFP and mCherry were excited by 488- and 561-nm laser lines respectively. Approximately 10-min sequential image acquisition was performed for microtubule and plasma-membrane time-lapses and 0.5-or-1 hourly for plasma-membrane and

PDLP time-lapses using an Evolve 512 camera (Photometrics) through band-pass filters (482/35 and 617/73; Semrock). Segment length as a function of segment's center of mass was conducted by first saving the XY-coordinates for both initial and final timepoints in separate '.csv' files with x and y headings then running the 'segmentLengthAFOCenterOfMass.py' code. Segments with at least 1 PDLP were identified and the 3-way-junctions were manually tagged using a 3-by-3 pixel square 1.5µm below the 3-way-junction for 2 image slices. Using the plasma-membrane, cell boundaries spanning from slightly passed the three-way junction were segmented manually using the segmented line tool with the spline function activated in FIJI and saved as an ROI. After all time-points were collected the ROI series were saved as an ".zip" file. The macro PDLP.ijm, which acts on both the plasma-membrane and PDLP channels at each time-point, straightens the segment, reslices the straightened segment, and then a max-projected image is produced. To reduce the tilt due to sample or mounting, one time-point was aligned to the x-axis using the 3-way-junction marks and the rest of the time-points were aligned to it using the resliceAlignment.ijm that worked on lines drawn from 3-way-junction to 3-way-junction. PDLPs were tracked using the Particle Tracker 2D/3D tool from the MOSAIC imaging toolset[16]. Non-overlapping PDLPs at least 2µm apart were used to calculate pair-wise distance as a function of time. These values were used to obtain a linear fit equation and its slope was used as the subsegmental growth rate; if the total displacement was less than image resolution then a growth rate of zero was awarded. To first verify the PDLP movement reflected growth, the displacement of 14 PDLP-pairs from 3 segments from 4 ˚C growth inhibited seedlings were compared to the growth rate from 38 PLDP-pairs from 12 segments from seedlings grown in 22 ˚C seedlings. To verify the accuracy of the PDLP sub-segmental growth, the sub-segment values were summed and evaluated to the displacement measured independently from the two endpoints defined by the 3-way-juncions. If the value was less than 3% error, then the measurements were accepted. Sub-segmental population level analysis was conducted as follows; sub-segments that occupied at least 60% of the lobe apex were classified as 'apex', similarly, sub-segments that occupied at least 60% of the lobe flank were classified as 'flank', everything else was grouped into the 'mix' sub-group. All statistical analyses were conducted using RStudio. Subsegmental growth rates plots were created in MATLAB. Python was run using Spyder IDE.

**Imaging of delamination and furrows after pectinase treatment and TEM microscopy.**
The plasma membrane marker PIP2:mCherry was excited by the 561nm laser line through bandpass filters described above. For low-light chemical treatment, 6-8 image fields at the basal region of the cotyledon were collected using the 63x C-Apo 1.2 NA water immersion objective and images were acquired using the Evolve 512 camera. For submerged chemical treatment, 1 DAG seedlings place in the chemical treatment for 24 hours, the then 2 DAG cotyledons were imaged completely using the 20x Plan-APO 0.8 NA objective using a Prime 95B camera (Photometrics), 3D montaging was done using the montage method in Slidebook. For TEM, leaf material was prepared as previously described[15].

**Finite element model**
Stress analysis in the pavement cell walls was studied using commercial finite element (FE) software Abaqus. The structural model of the cells was based on the segmented cell wall boundary and surrounding anticlinal wall. All cells were divided by a middle lamellar layer of pectin. The model before application of turgor pressure consisted of a flat periclinal wall bonded to the anticlinal walls. The thickness of the periclinal and anticlinal walls was 300 and 35 nm,

respectively. All cells were surrounded with pectin to approximate the confinement of adjacent cells. The turgor pressure in each cell was 0.6 MPa. The material for all cells was assumed to be a neo-Hookean, hyperelastic isotropic material assigned uniformly across the model. The elastic modulus was assumed as 600 MPa with a Poisson's ratio of 0.47. The whole material was assumed to be a standard linear solid with a primary relaxation time of 6.8 s with a ratio of infinite modulus to the total elastic modulus of 0.85[17]. The middle lamellar pectin was assumed to have the same properties as the surrounding pectin with an elastic modulus of 100 MPa[18]. For all pairs of segmented cells, after 20 sec of pressurization and sufficient relaxation, the stress field in the cell wall was used for analysis. For the segment of cells which were to study the correlation of segmental growth rate and the associated maximum principal stress, after the first step of 20 sec pressurization, the deformed structural model at the end of the first step was used as the initial structure in the second step and was pressurized for another 20 sec, and then the stress field in the walls at the end of second step was used for analysis.
To demonstrate the influence of the concentrated microfibrils (MF) located in the cell connection boundary for the lobe formation, one pair of pavement cells was used for finite element (FE) analysis. Considering the lobe formation during cell growth, the two cells were not confined using surrounding pectin, instead, pressure of 0.6 MPa were exerted against uniformly across the outer side of the whole anticlinal walls. Turgor pressure in the cells was also set to 0.6 MPa. The inner periclinal walls were constrained in the vertical direction given the support by the beneath cells, and one local region (random position) of the anticlinal walls were constrained in the lateral directions to ensure the complete boundary conditions of the system. Three steps were set for the analysis. The periclinal wall in the zero step was flat before pressurization and the inflated cell walls at the end of previous step were used as the initial structural models of the next step. The loading time at each step was set to 30 s for sufficient relaxation. At the zero step, all the materials were assumed to be isotropic neo-Hookean hyperelastic material with Young's modulus of 600 MPa and Poisson's ratio of 0.47, and they were assumed to be a standard linear solid with a primary relaxation time of 6.8 s having a ratio of infinite modulus to the total modulus of 0.85. In the following first and second steps, for the initiating cell, a patch of anisotropic materials, resembling a bundle of MF, was assigned for local periclinal and the associated anticlinal walls in the interaction region, and it has a rectangular area of 2x4 μm width and length in the periclinal wall. Local orthogonal material coordinates were defined for these materials in the region, with the x-axis perpendicular to the boundary edge and y-axis parallel with the edge. The materials in this region were assumed to be transversely isotropy and have mechanical response prescribed by Hooke's law. The parameters of the constitutive equation include elastic moduli of $E_{MF}$ = 2400 MPa and $E_T$ = 240 MPa, shear modulus of $G_{MF\_T}$ = 240 MPa, and Poisson's ratios of $\nu_{MF\_T}$ = 0.4 and $\nu_{T\_T}$ = 0.47. The modulus $E_{MF}$ was set in the x-axis, and $E_T$ was defined along the y and z-axis. A reference FE example was made to compare with the effect of presence of anisotropic patch. In the reference example, all the cell walls across the three steps were assumed to be isotropic neo-Hookean hyperelastic material. At the end of the final step, the connection boundary in both the reference example and the model using anisotropic patch was used for comparison.

**Anticlinal wall tilt analysis**

Segment shapes were traced as explained above. Detection of lobes and their location were identified by using the find_peaks, peaks_prominences modules in Python restricted to features greater than 286 nm as previously described[11]. To get the local tilt at the apex as a function of lobe formation, the relative position of the lobe in a normalized length was used to get its relative position prior to the lobe detection. A re-slice of the anticlinal wall at the apex was traced using the segmented line tool in Fiji for a time-point prior to lobe detection, at detection, and at least one time-point after detection. The traces were analyzed by first rotating them to the y-axis then measuring its width giving a score of zero to widths smaller than the imaging resolution. The OrientationJ Dominant Direction plug-in was used to obtain the tilt along the segment[19].

**Microtubule imaging time interval selection**
Single image and image stack time-lapses were first run through the background reduction as previously described[11]. Image stacks were max-projected. Timelapses were registered using rigid registration in StackReg using MT channel. Regions of interest were selected and cropped. Each timepoint was saved as a text image. The python code 'normalizedTextImageDataAndCorrelation.py' was run which first normalizes each image from 0 to 1. Then spearman correlations were conducted between, initial timepoint vs all timepoints, a moving window comparing a timepoint with the next timepoint, and finally between each timepoint and a randomly selected timepoint. The mean spearman correlation value for all columns was obtained for each comparison type listed above as a function of time. Values were then plotted. Statistical analysis and plots were done in Python using Spyder IDE.

**Microtubule persistence at the anticlinal and outer-periclinal wall for neighboring cells**
Using the segment traces generated as described above. The zip file for a segment's time-series was run through the imageTextOfMTWalls.ijm macro which allows the extraction of the microtubule signal from the anticlinal and outer-periclinal wall. For each time-point, the segment was straightened, re-sliced, and a SUM projected image was created of the plasma-membrane which allows the user to create a bounding box that defines the extent of the anticlinal wall on the image stack. This information was propagated to the microtubule channel. A text image of the outer-periclinal wall for each neighboring cell after a MAX intensity projection was produced. A text image of the anticlinal wall after SUM projection of 6 re-sliced image is also. produced. To create the persistence peaks of microtubules for all 3 cell wall views, the python code; createAnticlinalAndOuterPericlinalPersistenceMaps.py was run. This requires information about which cell is the initiating cell, what time-point the lobe is detected, the location of the lobe at detection, and the width of the lobe apex at detection. For each time-point, using the three matrices produced by imageTextOfMTWalls.ijm, the columns were summed to obtain a line plot for each of the cell wall views. The values for the anticlinal wall were normalized from zero to one using the equation: $normValue = \frac{Value_i - \min(array)}{\max(array) - \min(array)}$
For the periclinal wall, the values were normalized from zero to one using the maxima and minima of both datasets combined. Stacks-plots were created using the normalized value for the timepoints before lobe initiation, after lobe initiation, and for the whole time-lapse. MT enrichment was calculated by obtaining the area under the curve, using the trapezoid method, of the accumulated MT plots at the lobe apex. The enrichment is calculated using the following equation: $Enrichment = -1 + \frac{Area_{Cell1}}{Area_{Cell2}} * 100$
This is also done at each time-point individually to obtain the enrichment at each time-point.

**Microtubule orientation and coherency analysis**

Segments of interest were aligned to the first time-point by drawing a line across it's the 3-way-junctions and saving them as ROIs. The regIndSegWith3WJLines.ijm macro is then run which requires to select the location for the folder containing the max-projected images, the ROI zip file location from 3-way-junction lines, and an output folder. The location of a new lobe is then identified as explained above and two 2-by-2μm ROIs were placed across the anticlinal wall at the location where a new lobe will form. The OrientationJ Dominant Direction plug-in was used to obtain the orientation and coherency of microtubules. Plots were created in Python using Spyder IDE.

**Manual transfacial microtubule scoring**

Text image outputs from imageTextOfMTWalls.ijm were imported into Fiji and converted to RGB color. In order to observe which bundles were transfacial, it was necessary to combine the anticlinal and outer periclinal wall images using the combine tool in Fiji. The images were combined vertically with the anticlinal wall positioned directly below the outer periclinal wall that corresponded to the same time point. This was done for both cells that shared the anticlinal wall. Transfacial microtubule bundles were then scored by marking them in the middle of the bundle. Only clear, distinct transfacial microtubule bundles were scored, which we defined as microtubules that directly connected from the anticlinal wall into the outer periclinal wall. These bundles appear as a continuous, vertical line of high intensity pixels in the combined image. The pixel locations for all marked transfacial microtubule bundles, along with the pixel width of the entire image, were recorded in order to generate normalized segment length histograms. A density curve was obtained using these values. To match the y-value range to the histogram, the density values were multiplied by the proportional difference between the max value of the density plot and the maximal value of the histogram graph. Plots were created in RStudio.

**Microtubule organization within furrows and furrow position on straight segments**

Nascent furrows, obtained using the methods described above, were traced to and from regions where the anticlinal wall separated. For each furrow, the microtubule intensity was measured at the anticlinal wall and then normalized the signal from 0 to 1. The length of each furrow was normalized from 0 to 1. A stack-plot from 36 nascent furrows from 5 cotyledons was constructed from of the normalized signal to obtain a population level plot of microtubule preference within a furrow. The location of a furrow within a segment was obtained from the normalized peak location of a subset of nascent furrows from straight segments, 23 segments from 5 cotyledons, and a histogram was created using RStudio. Plots were created in Python using Spyder IDE.

**Local isotropic patch FEM simulation.**

To demonstrate the influence of the concentrated microfibrils (MF) located in the cell connection boundary for the lobe formation, one pair of pavement cells was used for finite element (FE) analysis. Considering the lobe formation during cell growth, the two cells were not constrained using surrounding pectin, instead, pressure of 0.6 MPa were exerted against uniformly across the outer side of the whole anticlinal walls. Turgor pressure in the cells was also set to 0.6 MPa. The inner periclinal walls were constrained in the vertical direction given the support by the beneath cells, and one local region (random position) of the anticlinal walls were constrained in the lateral directions to ensure the complete boundary conditions of the system. Three steps were set for the analysis. The periclinal wall in the zero step was flat before pressurization and the inflated cell walls at the end of previous step were used as the initial structural models of the next step. The loading time at each step was set to 30 s for sufficient relaxation. At the zero step, all the materials were assumed to be isotropic neo-Hookean hyperelastic material with Young's modulus of 600 MPa and Poisson's ratio of 0.47, and they were assumed to be a standard linear solid with a primary relaxation time of 6.8 s having a ratio of infinite modulus to the total modulus of 0.85. In the following first and second steps, for the initiating cell, a patch of anisotropic materials, resembling a bundle of MF, was assigned for local periclinal and the associated anticlinal walls in the interaction region, and it has a rectangular area of 2x4 μm width and length in the periclinal wall. Local orthogonal material coordinates were defined for these materials in the region, with the x-axis perpendicular to the boundary edge and y-axis parallel with the edge. The materials in this region were assumed to be transversely isotropy and have mechanical response prescribed by the Hooke's law. The parameters of the constitutive equation include elastic moduli of $E_{MF}$ = 2400 MPa and $E_T$ = 240 MPa, shear modulus of $G_{MF\_T}$ = 240 MPa, and Poisson's ratios of $\nu_{MF\_T}$ = 0.4 and $\nu_{T\_T}$ = 0.47. The modulus $E_{MF}$ was set in the x-axis, and $E_T$ was defined along the y and z-axis. A reference FE example was made to compare with the effect of presence of anisotropic patch. In the reference example, all the cell walls across the three steps were assumed to be isotropic neo-Hookean hyperelastic material. At the end of the final step, the connection boundary in both the reference example and the model using anisotropic patch was used for comparison.

**Code availability.**
FIJI macros, python scripts, and PLDP growth rates code in R are available at https://github.com/yamsissamy/naturePlant_lobeInitiation
